# Cellulose Synthase Complexes and Remorins Mediate Stress Resilience Through Cell Wall-Plasma Membrane Attachments

**DOI:** 10.1101/2025.08.01.664786

**Authors:** Yue Rui, Magda Zaoralová, William P. Dwyer, Andres V. Reyes, Tarabryn S. Grismer, Nikolaj B. Abel, Dharanidaran Jayachandran, Shishir P. S. Chundawat, Thomas Ott, Joseph J. Kieber, Peter D. Dahlberg, Shou-Ling Xu, José R. Dinneny

## Abstract

**Highlights:**

- The plasma membrane forms attachments to the plant cell wall that are revealed by hyperosmotic shock and correlate with tolerance to stress.
- Cellulose Synthase Complex (CSC) clusters and REMORIN (REM) nanodomains localize to cell wall-plasma membrane attachment sites.
- CSC density at the plasma membrane determines the extent of cell wall-plasma membrane attachment under hyperosmotic stress.
- REMs rapidly form nanodomains under hyperosmotic stress and are associated with SHOU4/4L, CSC exocytosis inhibitors that limit CSC density at the plasma membrane.

The outer cell surface of an organism is the frontline for detecting and responding to environmental stimuli. In plants, this interface consists of the plasma membrane that lies beneath the cell wall and remains associated with it through attachment sites. These wall-membrane attachments become evident upon hyperosmotic shock, when severe water loss causes the membrane to retract from the wall. Despite their long-standing observation, the molecular identity and function of these attachments remain poorly understood. Here, we identified two mechanisms governing wall-membrane attachments: one dependent on the Cellulose Synthase Complex (CSC), whose density at the plasma membrane positively correlates with resistance to hyperosmotic stress, and the other on REMORIN (REM), which acts antagonistically to the CSC mechanism. Using proximity-labeling proteomics, we identified SHOU4/4L as REM-associated proteins that mediate this antagonism. Together, our findings reveal how wall-membrane attachments are patterned to mediate plant cell resilience under water stress.

## Introduction

Living organisms must sense and respond to changing environments for their survival. The outer cell surface represents the frontline in this process and comprises an extracellular matrix that surrounds the cell and interfaces with the plasma membrane, where signaling is integrated and communicated to the cell through receptors and other membrane-associated proteins. A fundamental understanding of the organization, dynamics, and composition of this outer cell surface will shed light on the molecular underpinnings of stress perception and acclimation. In animal cells, the plasma membrane is anchored to the extracellular matrix by integrins,^1^ transmembrane receptors that serve as structural and signaling hubs for sensing and transmitting mechanochemical cues at the cell surface.^2^ Integrins have not been found in plants,^3,4^ however, and it remains unclear what molecules in the cell wall and the plasma membrane mediate their attachment, how the cell wall-plasma membrane interface is maintained and remodeled during environmental stress, and how signals are perceived at this interface in response to changes in internal (e.g., turgor pressure) or external cues (e.g., chemical or mechanical stimuli).

The plant cell wall is an extracellular hydrogel where cellulose microfibrils are embedded within a molecular matrix composed of hemicelluloses, pectins, and structural proteins.^5^ Plant cell walls play key roles in defining cell shape, providing structural support, and mediating cellular signaling.^6,7^ The cell wall constrains the lateral diffusion of plasma membrane proteins^8^ and regulates the size and dynamics of membrane nanodomains,^9^ which are submicron molecular assemblies with diverse protein and lipid compositions.^10,11^ Much of our understanding of cell wall function has come from studies in *Arabidopsis thaliana*, particularly through the characterization of mutants impaired in the synthesis, modification, or degradation of specific wall components.^5^ A number of studies have demonstrated the effects that environmental stress has on cell wall properties, and the physiological impact that disruptions in cell wall composition have on the ability of plants to acclimate to such stresses.^7,12^ However, whether specific wall constituents play a role in mediating wall-membrane attachments has not been determined.

Under non-stress conditions, turgor pressure appresses the plasma membrane against the cell wall (Figure 1, Figure S1A). However, under hyperosmotic stress, cellular water loss can be so severe that plasmolysis ensues, when the plasma membrane retracts from the cell wall (Figure 1C-1E, Figure S1A, Videos S1 and S2). Notably, during plasmolysis, portions of the plasma membrane remain attached to the cell wall, forming thin connecting membrane filaments known as Hechtian strands, first described by Hecht over a century ago.^13^ These strands are part of a larger network collectively referred to as the Hechtian structure,^14^ which includes Hechtian strands, the Hechtian reticulum, and wall-membrane attachment sites. The attachment sites, located at the distal ends of Hechtian structures (Figure 1D and 1E, Figure S1A, Videos S1 and S2) are reminiscent of focal adhesions in mammalian cells,^15^ yet their molecular identity and function remain largely unknown.^3,4^ Some Hechtian structures are likely attached to the wall via plasmodesmata,^14^ membrane-lined intercellular channels that mediate transport and communication between plant cells. However, the presence of wall-membrane attachment sites at the outer surface of epidermal cells^3^ and in guard cells,^16^ both of which lack plasmodesmata, suggests alternative mechanisms for connecting the cell wall and the plasma membrane.

**Figure 1.**
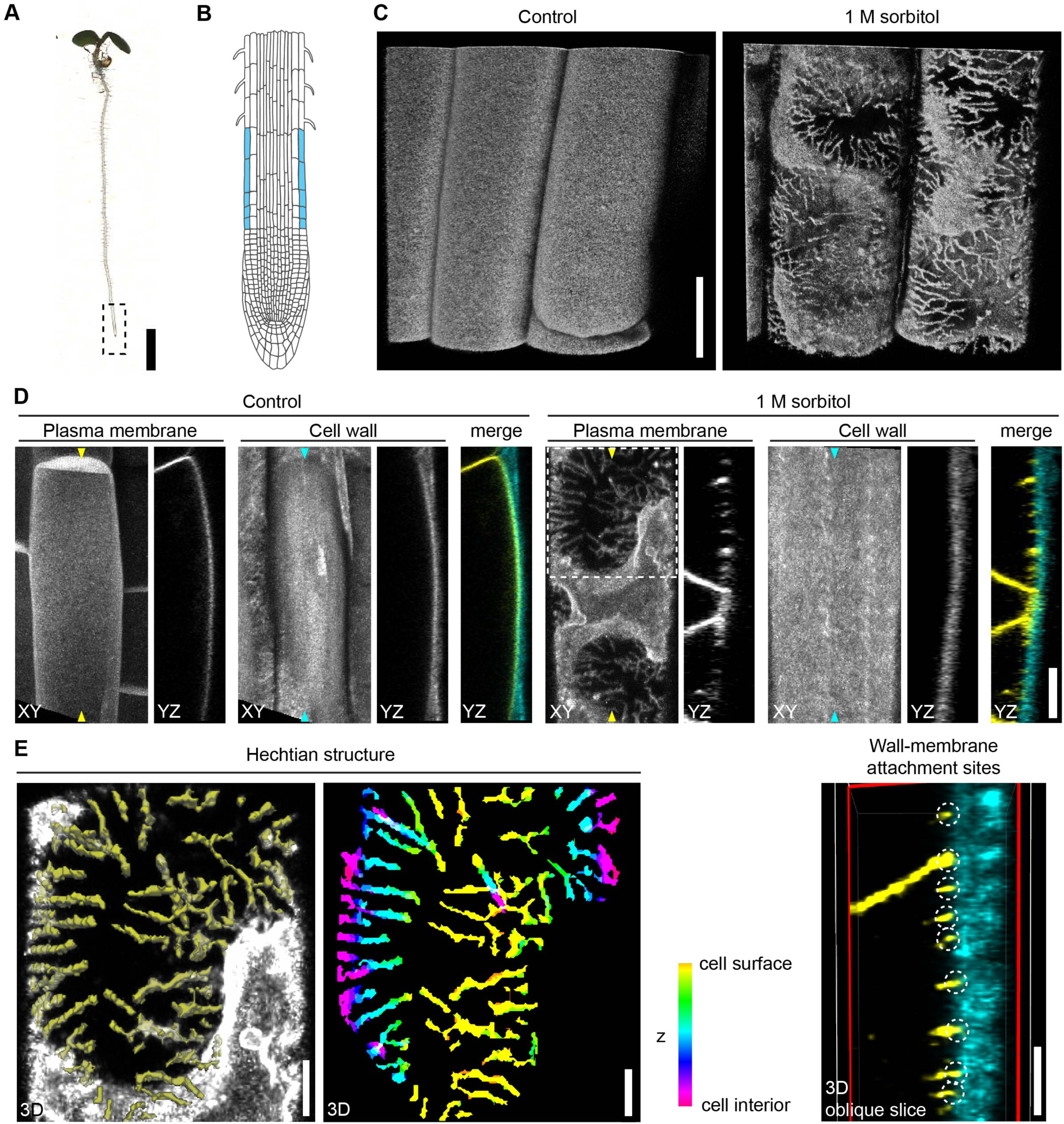
Hyperosmotic stress reveals cell wall-plasma membrane attachment sites in Arabidopsis roots. (A) A 5-d-old Arabidopsis seedling grown on MS media. Box demarcates the root tip. Scale bar: 0.2 cm. (B) Diagram of the longitudinal section of an Arabidopsis root tip in (A). Epidermal cells in the elongation zone are color coded in blue, which represent the regions where images in (C) were taken. (C) 3D rendering of root epidermal cells in 5-d-old seedlings expressing a plasma membrane marker, YFP-LTI6b, under the control condition or treated with 1 M sorbitol for 5 min. Scale bar: 10 µm. (D) Laser scanning confocal images of root epidermal cells from 5-day-old seedlings expressing the plasma membrane marker YFP-LTI6b and co-stained with the cell wall dye calcofluor white. Seedlings were imaged under control conditions or after treatment with 1 M sorbitol for 5 min. XY and YZ projections are shown. The positions where YZ projections were made are indicated by arrowheads. Scale bar: 5 µm. (E) 3D segmentation of Hechtian structure and wall-membrane attachment sites from the dashed box region in (D). Segmented Hechtian structure is shown either in yellow, overlaid on YFP-LTI6b signals, or with depth-based color coding. Wall-membrane attachment sites (indicated by white dashed circles) are visualized by applying an oblique slicer (outlined in red) to the YZ projection. Scale bars: 2 µm. See also Video S1.

The plasma membrane is a lipid bilayer enriched with diverse lipid species and proteins, and it exhibits heterogeneity in its physicochemical properties.^17^ Importantly, many membrane proteins are organized into clusters or nanodomains to fulfill their functions in plant growth, abiotic stress responses, and biotic interactions.^10,11^ For example, Cellulose Synthase Complexes (CSCs) appear as discrete puncta at the plasma membrane when imaged by confocal microscopy.^18^ CSCs have been proposed to anchor the plasma membrane to the cell wall in plasmolyzed plant cells through the nascently synthesized cellulose microfibrils that become embedded into the wall matrix,^19^ yet this idea remains experimentally untested. Hyperosmotic stress can also induce the formation of nanodomains, which was recently demonstrated for ROP6, a Rho GTPase that interacts with NADP oxidases to produce reactive oxygen species important for osmotic signaling.^20^ In the context of biotic interactions, REMORINs (REMs) are a well-characterized protein family known to organize into nanodomains.^21^ For instance, the symbiosis-specific remorin SYMREM1 in *Medicago truncatula* is essential for stabilizing the symbiotic receptor-like kinase LYK3^22^ and maintaining membrane topology during infection.^23^ Interestingly, *in vitro* studies have shown that REMs can bind oligo- and polygalacturonides,^24^ components of pectins in the cell wall. However, whether REMs contribute to physical or functional associations between the cell wall and plasma membrane is unknown.

Plant roots absorb water and nutrients from the environment, and their growth is highly sensitive to environmental fluctuations. Here, we leveraged the effects of hyperosmotic stress to investigate the structure and function of the cell wall-plasma membrane interface in roots. Our analyses focused on epidermal cells in the elongation zone, where rapid cell expansion places a strong demand on the integrity of the wall-membrane interface.^25^ Root hair cells were excluded because they are highly susceptible to hyperosmotic stress-induced damage and frequently undergo cell death under these conditions. In addition, epidermal cells in the elongation zone are readily accessible for high-resolution live-cell imaging, enabling quantitative analysis of cell wall-plasma membrane organization during osmotic stress. We show that wall-membrane attachment sites are marked by clusters of CSCs and REM nanodomains. Maintaining CSC density at the plasma membrane is essential for wall-membrane attachments and sustained root growth under hyperosmotic stress conditions. Genetic, biochemical, and co-localization studies identified SHOU4/4L, known negative regulators of CSC exocytosis,^26^ as being associated with REMs that limit CSC density at the plasma membrane. Our data support a one-way antagonistic mechanism where REMs restrict the abundance of CSCs. CSCs and REMs function together at the plasma membrane to regulate wall-membrane attachments, which are critical for maintaining cellular resilience under water deficit conditions.

## Results

### Impaired cellulose and rhamnose levels have contrasting effects on root growth under hyperosmotic stress

To investigate the roles of cell wall polysaccharides in hyperosmotic stress response, we surveyed a collection of previously published Arabidopsis mutants deficient in cellulose, xyloglucan (a hemicellulose), pectic homogalacturonan (HG), rhamnogalacturonan-I (RG-I), or rhamnogalacturonan-II (RG-II), respectively, for changes in root growth (Figure S1B, Figure 2). Under hyperosmotic stress induced by 0.28 M sorbitol, wild-type roots exhibited a growth reduction of 52-57% compared to the control conditions (Figure 2). Mutants lacking cellulose, including *cesa3*^je5^, *cesa6^prc^*^1–1^, and *cob-1*, all displayed a more severe growth reduction (≥75%) (Figure 2A). Expression of *GFP-CESA3* under its native promoter in the *cesa3^je^*^5^ mutant background restored root growth to wild-type levels (Figure 2A). We selected *cesa3^je^*^5^ and *cesa6^prc^*^1–1^ for further growth rate analysis and found that their growth rate remained low and failed to recover over time under 0.28 M sorbitol conditions (Figure S1C and S1D).

**Figure 2.**
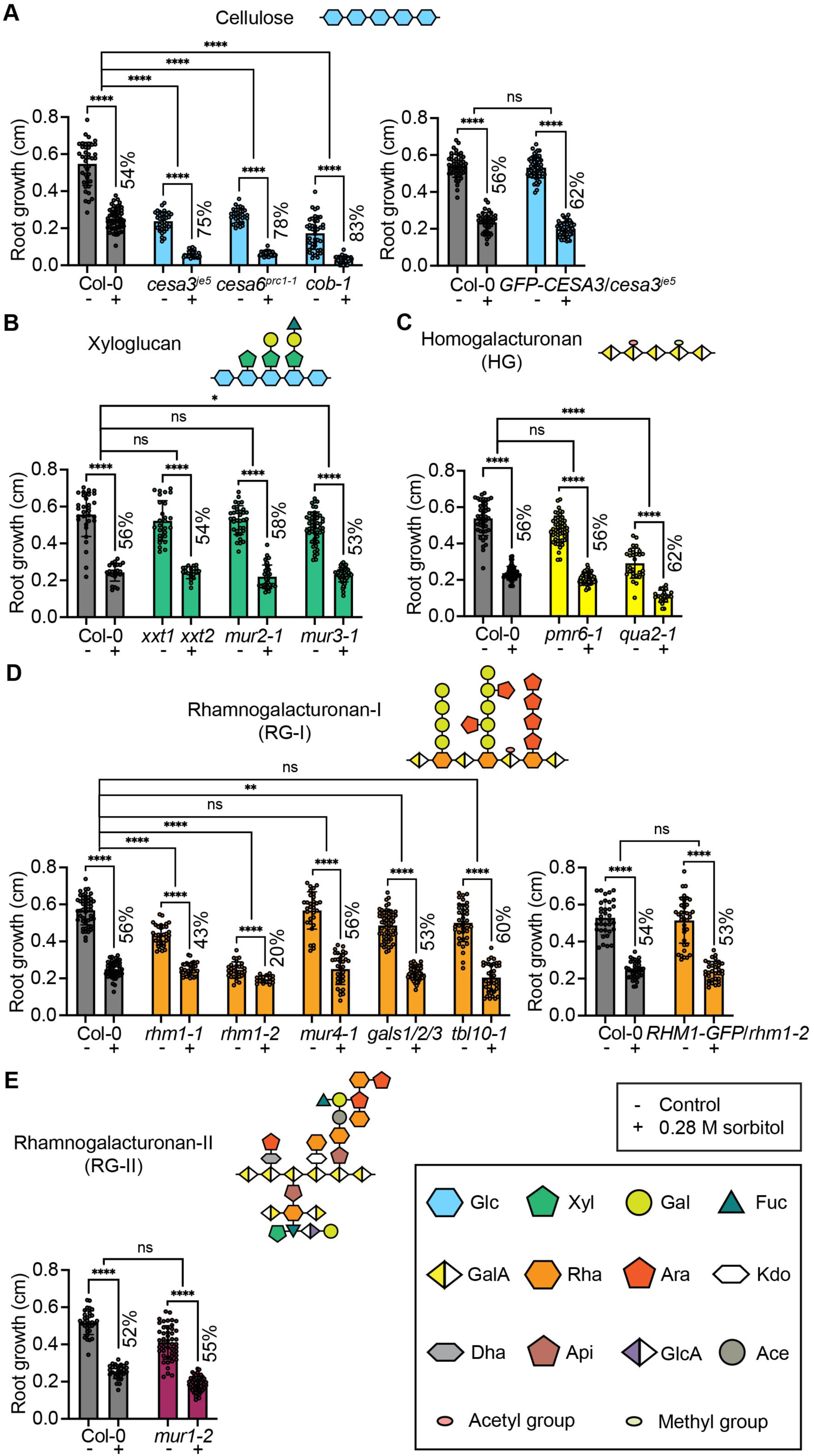
Reductions in cellulose and rhamnose impact root growth under hyperosmotic stress in opposing ways. (A) Root growth of wild type, cellulose-deficient mutants, and a *CESA3* complementation line 1 day after transfer to either fresh MS media (-) or MS media supplemented with 0.28 M sorbitol (+). *n* ≥ 27 seedlings per genotype per treatment. (B) Root growth of wild type and xyloglucan mutants 1 day after transfer to either fresh MS media (-) or MS media supplemented with 0.28 M sorbitol (+). *n* ≥ 20 seedlings per genotype per treatment. (C) Root growth of wild type and homogalacturonan mutants 1 day after transfer to either fresh MS media (-) or MS media supplemented with 0.28 M sorbitol (+). *n* ≥ 25 seedlings per genotype per treatment. (D) Root growth of wild type and rhamnogalacturonan-I mutants 1 day after transfer to either fresh MS media (-) or MS media supplemented with 0.28 M sorbitol (+). *n* ≥ 16 seedlings per genotype per treatment. (E) Root growth of wild type and rhamnogalacturonan-II mutants 1 day after transfer to either fresh MS media (-) or MS media supplemented with 0.28 M sorbitol (+). *n* ≥ 29 seedlings per genotype per treatment. Error bars indicate SD. ns, no significance; \**p* < 0.05, \*\**p* < 0.01, \*\*\**p* < 0.001, \*\*\*\**p* < 0.0001 by two-way ANOVA.

Mutants defective in xyloglucan (Figure 2B), HG (Figure 2C), RG-I acetylation (Figure 2D), or RG-II crosslinking (Figure 2E) generally showed wild-type responses to stress. The only exception was *qua2-1*, which exhibited a significantly greater reduction in root growth compared to wild type under stress (Figure 2C), though this may be influenced by its known defects in cell-cell adhesion.^27^

In contrast to the cellulose mutants, the *rhm1-1*, *rhm1-2*, and *rhm1-3* mutants, which are deficient in rhamnose, a component of the RG-I backbone,^28^ showed significantly less reduction in root growth under hyperosmotic stress than the wild type (Figure 2D and Figure S1E). This phenotype was reversed by expressing *RHM1-GFP* under its native promoter (Figure 2D).

Rhamnose in RG-I can be substituted with side chains, including *α*-1,5-arabinan and *β*-1,4-galactan. While a deficiency in galactan, as in the *gals1/2/3* mutants,^29^ partially mirrored the *rhm1* mutant’s response, a deficiency in arabinan, as in the *mur4-1* mutant, did not show a similar effect (Figure 2D). Taken together, our mutant survey suggests specific but contrasting roles of cellulose and RG-I rhamnose in root growth responses to hyperosmotic stress.

### Deficiency in cellulose or rhamnose has opposite effects on plasmolysis and baseline plasma membrane proteome

To further understand the functions of cellulose and rhamnose in hyperosmotic stress response at the cellular and molecular level, we performed live-cell imaging of five-day-old seedlings expressing YFP-LTI6b, a classic plasma-membrane marker, after a brief hyperosmotic stress treatment. The 0.28 M sorbitol concentration used in our root growth assay (Figure 2) did not cause any gross changes in overall root cell morphology compared to the control condition (Figure S1F). However, subtle deformations were consistently observed at cell corners, suggesting that cell shrinkage may initiate at these sites. To quantify this effect, we measured the angle formed between adjacent corners of two neighboring epidermal cells (Figure S1G). Compared with the wild type, cellulose-deficient *cesa3^je^*^5^ mutant cells displayed significantly wider corner angles, whereas rhamnose-deficient *rhm1-1* mutant cells exhibited markedly smaller angles (Figure S1G and S1H). By contrast, no significant differences were observed among genotypes under control conditions, in which corner angles were essentially 0° and therefore not meaningful to quantify (Figure S1G). These results reveal cell corner geometry as a sensitive feature in response to mild hyperosmotic stress. Treatment with 0.5 M sorbitol induced plasmolysis, with the plasma membrane detaching from the cell wall at cell-cell junctions along the root’s longitudinal axis (Figure S1F). Increasing the sorbitol concentration to 1 M caused a more pronounced plasmolysis effect, with the membrane also retracting from the outer periphery of root cells. Hechtian structures and cell wall-plasma membrane attachment sites were clearly visible at these membrane retraction sites (Figure 1C-1E, Figure S1A and S1F, Videos S1 and S2).

We selected 0.5 M sorbitol to compare plasmolysis in the wild type, *cesa3^je^*^5^, and *rhm1-1* mutants, as this concentration is less stressful than 1 M sorbitol and allows for easier quantification of plasmolysis by area. The ratio of protoplast area (highlighted by YFP-LTI6b) to cell area (outlined by the cell wall in the bright field channel) was used as a measure of plasmolysis. Compared to wild-type cells, plasmolysis was more severe in the *cesa3^je^*^5^ mutant, with a greater proportion of cells displaying lower ratios between 0.6-0.8 (26% in *cesa3^je^*^5^ vs. 12% in wild type) (Figure 3A and 3B). In contrast, 70% of the *rhm1-1* cells exhibited minimal plasmolysis at a ratio between 0.9-1.0, whereas this proportion in the wild type is 34% (Figure 3A and 3B). These cellular-level plasmolysis results correlate with the severity with which root growth was inhibited in these mutants (Figure 2A and 2D). Similar plasmolysis effects were observed in etiolated hypocotyls (Figure S2A), which also undergo rapid cell expansion, suggesting that alterations in cell wall composition consistently lead to differential plasmolysis responses across organ types.

**Figure 3.**
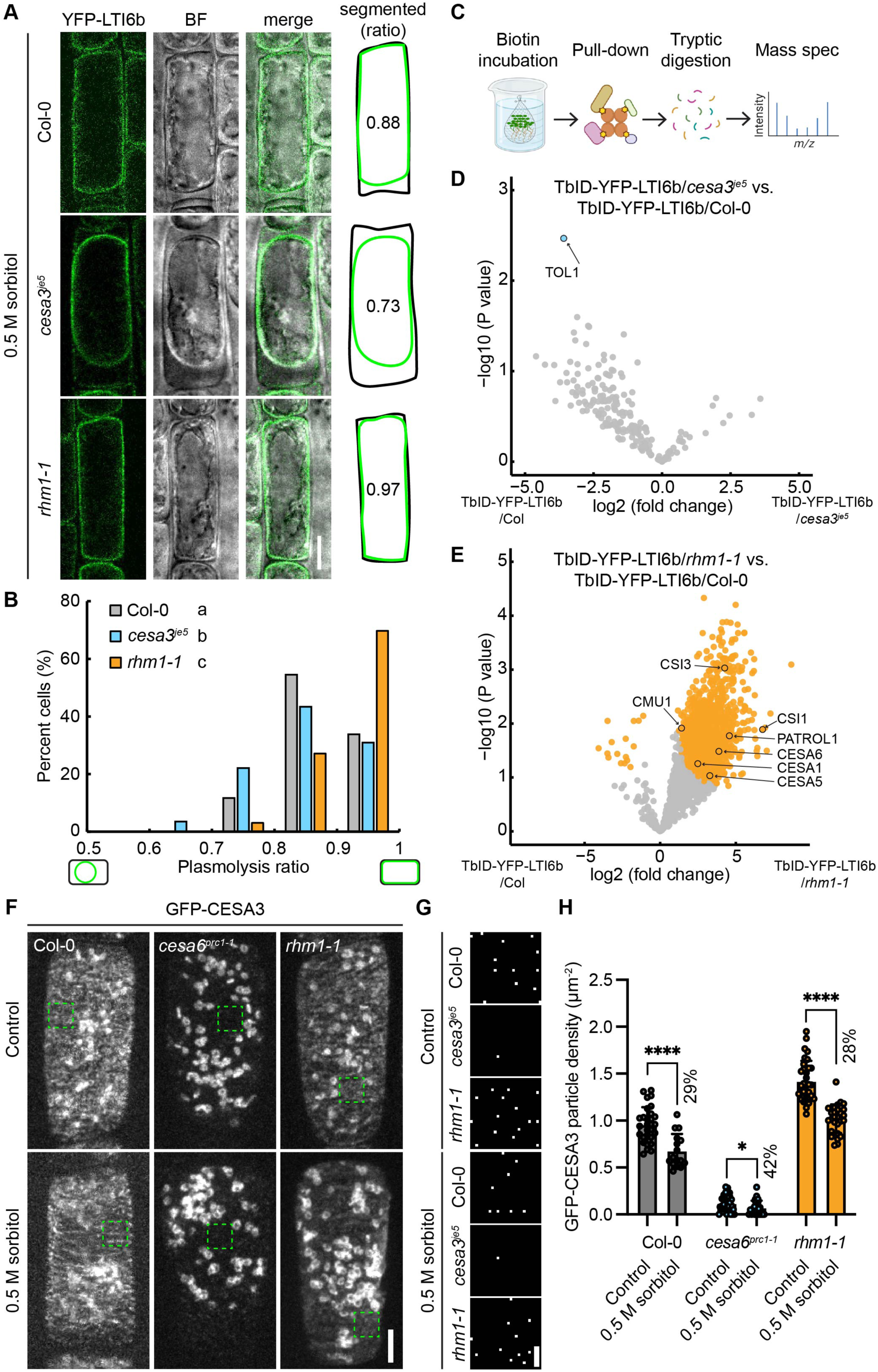
Deficiency in cellulose or rhamnose has opposite effects on plasmolysis and the plasma membrane proteome. (A) Laser scanning confocal images and manual tracings of plasmolyzed root epidermal cells in 5-d-old seedlings expressing YFP-LTI6b in Col-0, *cesa3^je^*^5^, or *rhm1-1* background, respectively. Plasmolysis was induced by 0.5 M sorbitol for 5 min. Each displayed ratio corresponds to the same individual cell. Scale bar: 10 µm. (B) Histogram showing plasmolysis ratio (protoplast area to cell area) in root epidermal cells of 5-d-old seedlings expressing YFP-LTI6b in Col-0, *cesa3^je^*^5^, or *rhm1-1* background, respectively. Different letters indicate significant differences by one-way ANOVA and Tukey’s test. *n* ≥ 113 cells from at least 10 seedlings per genotype. (C) Schematic workflow of the proximity labeling-based mass spectroscopy. (D) Volcano plot showing differentially enriched proteins in the plasma membrane proteome of *cesa3^je^*^5^ compared to the wild type Col-0. Significantly enriched or depleted proteins are color coded in blue. Pre-filtering steps removed genetic background from each TbID-YFP-LTI6b-labeled samples using Welch t-test with an FDR of 0.05 and an S0 of 0.5. Pair-wise comparison was then conducted between TbID-YFP-LTI6b/*cesa3^je^*^5^ and TbID-YFP-LTI6b/Col-0 using Student’s *t*-test with an FDR of 0.05 and an S0 of 0.5. *n* ≥ 3 biological replicates per genotype. (E) Volcano plot showing differentially enriched proteins in the plasma membrane proteome of *rhm1-1* compared to the wild type Col-0. Significantly enriched or depleted proteins are color coded in orange. Pre-filtering steps removed genetic background from each TbID-YFP-LTI6b-labeled samples using Welch t-test with an FDR of 0.05 and an S0 of 0.5. Pair-wise comparisons were then conducted between TbID-YFP-LTI6b/*rhm1-1* and TbID-YFP-LTI6b/Col-0 using Student’s *t*-test with an FDR of 0.05 and an S0 of 0.5. *n* ≥ 3 biological replicates per genotype. (F) Spinning disk confocal images of GFP-CESA3 in root epidermal cells of 5-d-old seedlings of Col-0, *cesa6^prc^*^1–1^, and *rhm1-1* under the control condition or induced by 0.5 M sorbitol for 5 min. Scale bar: 5 µm. (G) Magnified views of the regions outlined by green dashed boxes in (F), showing detected GFP-CESA3 particles. Scale bar: 1 µm. (H) Quantification of GFP-CESA3 particle density at the plasma membrane from (F). Error bars indicate SD. *N* ≥ 16 cells from at least 7 seedlings per genotype per treatment. \**p* < 0.05 and \*\*\*\**p* < 0.0001 by Student’s *t*-test.

We next determined whether the extent of plasmolysis relates directly with the ability of cells to recover growth from osmotic shock. While root cell elongation was reduced by 42% in the wild type during the first hour after a 0.5 M sorbitol shock, *cesa3^je^*^5^ mutants exhibited a greater reduction of 67% (Figure S2B and S2C). At the organ scale, overall root growth one day after a 0.5 M sorbitol shock was similarly reduced in *cesa3^je^*^5^ mutants but was significantly enhanced in *rhm1-1* mutants (Figure S2D). These data indicate that the ability to maintain wall-membrane attachment correlates well with the extent of recovery following osmotic shock.

To identify membrane protein signatures underlying the distinctive hyperosmotic stress responses of *cesa3^je^*^5^ and *rhm1-1* mutants, we used a biotin ligase-based proximity labeling approach^30,31^ by expressing a TurboID (TbID)-YFP-LTI6b fusion protein in the respective genetic backgrounds. After confirming that TbID-YFP-LTI6b localized to the plasma membrane (Figure S2E), we treated five-day-old seedlings with 50 µM biotin for 15 min (Figure 3C), which was sufficient to biotinylate proteins beyond the endogenously biotinylated ones, as confirmed by western blotting (Figure S2F and S2G). Following protein extraction, streptavidin pull-down of the biotinylated proteins, tryptic digestion, and mass spectrometry, we conducted pairwise protein enrichment analyses between each mutant and the wild type (Figure S2H and S2I).

In the *cesa3^je^*^5^ mutant, only one protein exhibited differential labeling: TOM1-LIKE1 (TOL1),^32,33^ which was significantly less enriched compared to wild type (Figure 3D, Table S1). In contrast, over a thousand proteins showed differential enrichment in the *rhm1-1* mutant relative to wild type (Figure 3E, Table S1). Among the most enriched proteins were CESA1, CESA5, CESA6, CELLULOSE SYNTHASE-INTERACTIVE1 (CSI1),^34,35^ CSI3,^36^ PATROL1,^37^ and CMU1^38^

(Figure 3E, Table S1), which are involved in cellulose synthesis, CSC delivery to the plasma membrane, or microtubule stability to guide CSC movement. These differences in enrichment were not caused by changes in LTI6b levels, which were consistent across all transgenic lines (Table S1). These data suggest that defects in RG-I rhamnose biosynthesis lead to an increase in the abundance of CSC-associated proteins at the plasma membrane.

To directly measure CSC density at the plasma membrane in root cells and assess its changes due to deficiencies in cellulose or rhamnose, we introduced the GFP-CESA3 marker, driven by its native promoter, into the wild-type, *cesa6^prc^*^1–1^, and *rhm1-1* backgrounds. Similar to earlier reports in etiolated hypocotyls,^18^ CSCs were observed at the plasma membrane as particles near or below the resolution limit of the spinning disk confocal microscope, and in Golgi as bright “disks” larger than the membrane-localized particles (Figure 3F). Compared to the wild type, GFP-CESA3-labeled CSC particles in *cesa6^prc^*^1–1^ roots were barely detectable at the plasma membrane, though Golgi-localized CSCs remained visible (Figure 3F-3H). In contrast, *rhm1-1* roots exhibited a significantly higher CSC particle density at the plasma membrane than wild-type roots (Figure 3F-3H), consistent with our proteomic characterization (Figure 3E).

These patterns of CSC density at the plasma membrane were maintained in seedlings treated with 0.28 M sorbitol stress (Figure S2J) and in seedlings subjected to an acute 0.5 M sorbitol shock (Figure 3F-3H). In the latter case, however, CSC density was reduced in all three genotypes relative to the control condition. Together, these results suggest that CSC density at the plasma membrane under normal conditions establishes the baseline for CSC density under hyperosmotic stress.

### CSCs mediate cell wall-plasma membrane attachments under hyperosmotic stress

Our results thus far point to a possible link between CSC density at the plasma membrane and the severity of plasmolysis during hyperosmotic stress, suggesting that CSC may be directly or indirectly involved in promoting wall-membrane attachments. To test this hypothesis, we first visualized CSCs at the plasma membrane and the cell wall using the GFP-CESA3 marker co-stained with Alexa568-tdCBM3a, a probe specific for cellulose.^39^ CSCs were observed to associate with cellulose under both control conditions and following 1 M sorbitol treatment (Figure S3A). In addition, we crossed the tdTomato-CESA6 marker with GFP-LTI6b and imaged root cells using spinning disk confocal microscopy. Under control conditions, tdTomato-CESA6-labeled CSCs appeared as particles that move along linear trajectories at the plasma membrane (Figure 4A, Figure S3B, Video S3), consistent with previous observations in etiolated hypocotyls.^18^ Treatment with 1 M sorbitol immobilized CSCs into clusters at the plasma membrane (Figure S3B, Video S3), which showed an increased degree of signal overlap with GFP-LTI6b-labeled wall-membrane attachment sites compared to the control condition (Figure 4A and 4B, Video S4). To further investigate the formation of CSCs into clusters, we tracked their temporal dynamics at the plasma membrane before and after treatment with 1 M sorbitol (Figure S3C, Video S5). Some CSC clusters were already present immediately after hyperosmotic stress treatment, while others formed through the aggregation of multiple CSC particles (Figure S3C and S3D, Video S5). To complement time-lapse analyses of CSC spatial reorganization, we quantified CSC fluorescence intensity and particle density at the plasma membrane under control and 1 M sorbitol conditions. CSC intensity at the membrane increased approximately twofold following sorbitol treatment (Figure S3E), while CSC particle density was reduced relative to control conditions (Figure S3F; Figure 3H), with no significant difference between 0.5 M and 1 M sorbitol treatments (*p* = 0.16, 0.92, and 0.08 for Col-0, *cesa6^prc^*^1–1^, and *rhm1-1*, respectively). These observations indicate that hyperosmotic stress drives the redistribution of pre-existing CSCs into fewer, higher-intensity clusters, consistent with the aggregation events observed in time-lapse imaging. Nevertheless, under hyperosmotic stress, CSC particle density remained significantly higher in *rhm1-1* mutants and significantly lower in *cesa6^prc^*^1–1^ mutants compared with the wild type (Figure S3F).

**Figure 4.**
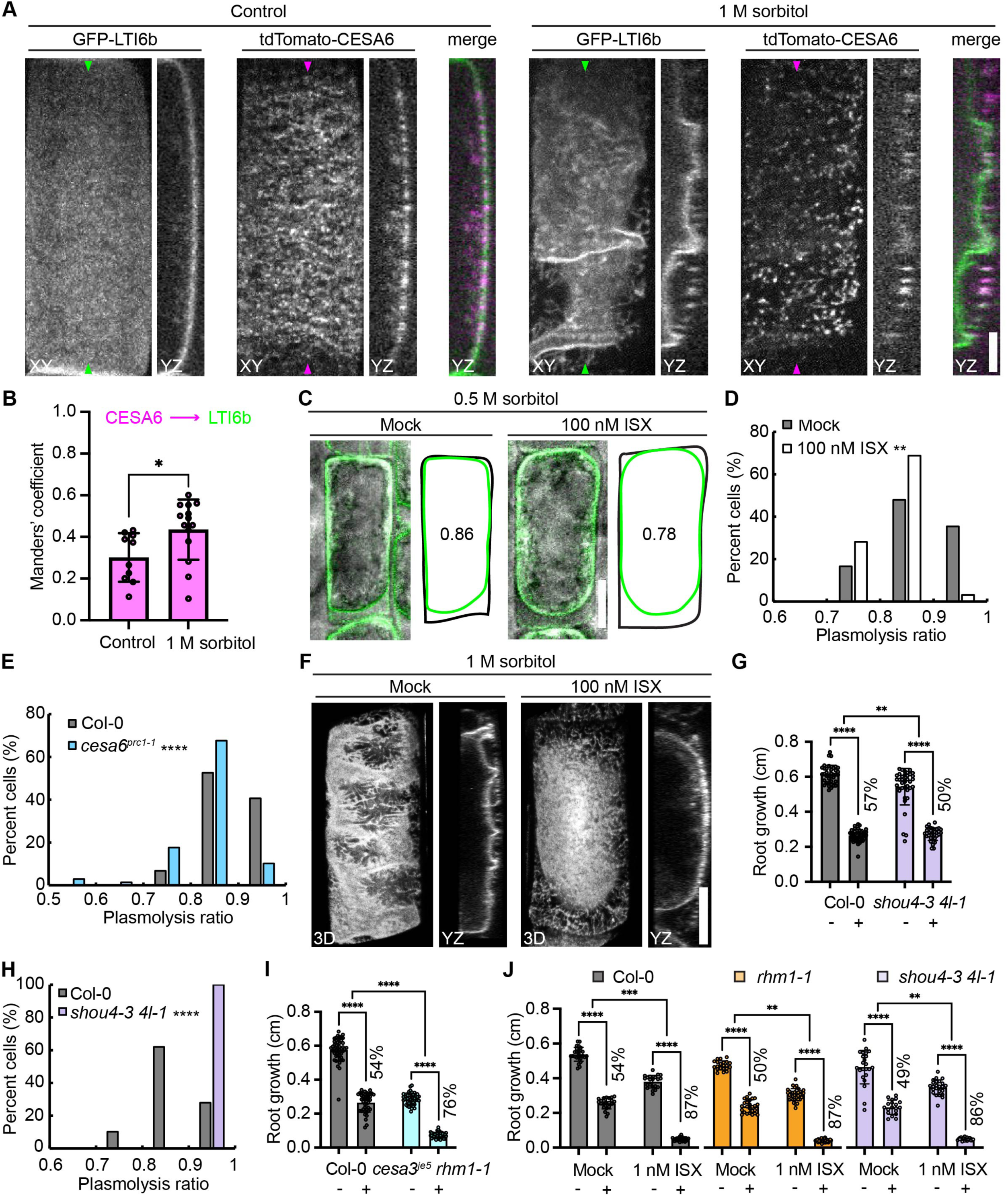
CSC density at the plasma membrane safeguards wall-membrane attachments and plant growth under hyperosmotic stress. (A) Spinning disk confocal images of root epidermal cells in 5-d-old seedlings expressing GFP-LTI6b and tdTomato-CESA6 under the control condition or treated with 1 M sorbitol for 5 min. XY and YZ projections are shown. The positions where YZ projections were made are indicated by arrowheads. Scale bar: 5 µm. (B) Quantification of Manders overlap coeffiicient in 5-d-old seedlings of GFP-LTI6b/tdTomato-CESA6 under the control condition or treated with 1 M sorbitol for 5 min. Error bars indicate SD. *n* ≥ 10 cells per genotype per treatment. \**p* < 0.05 by Student’s *t*-test. (C) Laser scanning confocal images and manual tracings of plasmolyzed root epidermal cells in 5-d-old Col-0 seedlings expressing YFP-LTI6b, treated with or without 100 nM isoxaben (ISX) for 20 minutes. Plasmolysis was subsequently induced by treatment with 0.5 M sorbitol for 5 minutes. Each displayed ratio corresponds to the same individual cell. Scale bar: 10 µm. (D) Histogram showing plasmolysis ratio in root epidermal cells in 5-d-old Col-0 seedlings expressing YFP-LTI6b, treated with or without 100 nM ISX for 20 minutes. *n* ≥ 32 cells from at least 5 seedlings per treatment. \*\**p* < 0.01 by Student’s *t*-test. (E) Histogram showing plasmolysis ratio induced by 0.5 M sorbitol in root epidermal cells of 5-d-old seedlings expressing YFP-LTI6b in Col-0 or *cesa6^prc^*^1–1^ background, respectively. *n* ≥ 59 cells from at least 10 seedlings per genotype. \*\*\*\**p* < 0.0001 by Student’s *t*-test. (F) 3D renderings and YZ projections of root epidermal cells of 5-d-old YFP-LTI6b seedlings treated with or without 100 nM ISX for 20 minutes. Plasmolysis was subsequently induced by treatment with 1 M sorbitol for 5 minutes. Scale bar: 10 µm. (G) Root growth of wild type and *shou4-3 4l-1* mutants 1 day after transfer to either fresh MS media (-) or MS media supplemented with 0.28 M sorbitol (+). *n* ≥ 34 seedlings per genotype per treatment. (H) Histogram showing plasmolysis ratio induced by 0.5 M sorbitol in root epidermal cells of 5-d-old seedlings expressing YFP-LTI6b in Col-0 or *shou4-3 4l-1* background, respectively. *n* ≥ 79 cells from at least 9 seedlings per genotype. \*\*\*\**p* < 0.0001 by Student’s *t*-test. (I) Root growth of wild type and *cesa3^je^*^5^ *rhm1-1* mutants 1 day after transfer to either fresh MS media (-) or MS media supplemented with 0.28 M sorbitol (+). *n* ≥ 42 seedlings per genotype per treatment. (J) Effects of ISX on root growth in wild type, *rhm1-1*, and *shou4-3 4l-1* mutants 1 day after transfer to either fresh MS media (-) or MS media supplemented with 0.28 M sorbitol (+) in the presence or absence of 1 nM ISX. *n* ≥ 19 seedlings per genotype per treatment. Error bars indicate SD. \*\**p* < 0.01, \*\*\**p* < 0.001, \*\*\*\**p* < 0.0001 by two-way ANOVA.

Because cortical microtubules guide the distribution of CSCs at the plasma membrane,^18,40^ we also examined how acute hyperosmotic stress affects microtubules and their colocalization with CSCs using the GFP-CESA3/mCherry-TUA5 double marker line.^40^ mCherry-TUA5-labeled microtubules appeared as strand-like structures that mirrored Hechtian structures, as previously reported,^41^ and were decorated by the immobilized population of CSCs at the plasma membrane (Figure S4A). In addition to this population of CSCs associated with microtubules, CSCs were also observed outside these regions (Figure S4A). Quantitative analysis shows that the density of CSCs is higher in regions of the cell periphery that remain associated with microtubules (Figure S4B).

To determine whether plasma membrane-localized CSCs mediate wall-membrane attachments, we treated YFP-LTI6b seedlings with the cellulose biosynthesis inhibitor isoxaben (ISX), which has previously been shown to remove CSCs from the plasma membrane.^18^ To establish conditions that produce an acute clearance of CSCs, we compared two ISX concentrations. Both 1 nM and 100 nM ISX significantly reduced CSC density in root cells; however, 100 nM ISX was more effective at achieving robust CSC removal within 20 min (Figure S4C-S4F). Based on this result, we treated seedlings with 100 nM ISX, induced plasmolysis with 0.5 M sorbitol and quantified the degree of plasmolysis (Figure 4C). Compared to the mock condition, ISX treatment resulted in a greater fraction of cells exhibiting increased plasmolysis (Figure 4D). This trend is consistent with that observed in the cellulose-deficient mutants *cesa3^je^*^5^ (Figure 3A and 3B) and *cesa6^prc^*^1–1^ (Figure 4E). Additionally, ISX caused changes in protoplast morphology under 0.5 M sorbitol, specifically enhancing curvature at both ends (Figure 4C, Figure S4G). To further assess the impact of ISX on wall-membrane attachments at the outer cell surface, we induced plasmolysis using 1 M sorbitol. In mock-treated roots, wall-membrane attachment sites were evident in regions where the plasma membrane had dissociated from the wall of the outer cell surface. In contrast, in ISX-treated roots, protoplast detachment and shrinkage occurred primarily at the cell-cell junctions (Figure 4F).

To determine the physiological relevance of disrupting wall-membrane attachments, we assessed its impact on growth recovery following hyperosmotic shock in ISX-treated seedlings. We used a lower concentration of ISX (1 nM) to reduce CSC density without compromising seedling viability. Compared with mock-treated controls, ISX-treated roots showed reduced cell elongation during the first hour after a 5-min 0.5 M sorbitol shock, although this difference was not statistically significant (Figure S4H). At the organ scale, ISX-treated seedlings exhibited significantly reduced root growth one day after a 30-min 0.5 M sorbitol shock (Figure S4I).

These results are consistent with the growth recovery defects observed in *cesa3^je^*^5^ mutants (Figure S2B-S2D) and support the hypothesis that wall-membrane attachment sites contain CSCs, which are necessary for the physical stability of these associations during hyperosmotic stress.

To further test this hypothesis, we analyzed another cell wall mutant, *shou4-3 4l-1*, which lacks the negative regulators of CSC exocytosis, SHOU4 and SHOU4L, and was previously shown to exhibit a higher CSC density at the plasma membrane relative to the wild type in etiolated hypocotyls.^26^ After confirming that CSC density was increased in *shou4-3 4l-1* root cells under both control conditions and 1 M sorbitol treatment (Figure S4J and S4K), we next examined their growth and plasmolysis phenotypes. Inhibition of root growth under hyperosmotic stress was significantly less severe in *shou4-3 4l-1* compared to the wild type (Figure 4G). Consistent with this observation, *shou4-3 4l-1* also showed improved root growth recovery following a 0.5 M sorbitol shock compared with the wild type (Figure S4L). Moreover, the proportion of root cells in this mutant that exhibit no measurable wall-membrane detachment was substantially higher than that in the wild type (Figure 4H), resembling the phenotypes observed in the *rhm1-1* mutants (Figure 2D, Figure 3A and 3B, Figure S2D). The *cesa6^prc^*^1–1^ *shou4-1* double mutant,^26^ however, displayed root growth phenotypes similar to the *cesa6^prc^*^1–1^ single mutant (Figure S4M, Figure 2A), suggesting that the *cesa6^prc^*^1–1^ mutation is epistatic to the *shou4-1* mutation.

Similarly, the *cesa3^je^*^5^ mutation was epistatic to *rhm1-1* in root growth under both control and hyperosmotic stress conditions (Figure 4I, Figure 2A). As a complementary approach, we tested the effect of ISX on root growth in the *rhm1-1* mutant or *shou4-3 4l-1* double mutant and found that this treatment similarly led to a more severe degree of growth reduction than the non-ISX treated controls (Figure 4J). These results indicate that CSCs act downstream of *RHM1* and *SHOU4/4L*.

Together, these data suggest that CSC density at the plasma membrane safeguards hyperosmotic stress responses in roots by mediating wall-membrane attachments.

### REMs rapidly form nanodomains at wall-membrane attachment sites upon hyperosmotic stress

CSCs are organized as discrete punctate structures at the plasma membrane under both control and hyperosmotic stress conditions (Figure 4A). To investigate how plasma membrane nanodomains, also appearing as puncta but defined as multi-component local compartments,^10^ respond to hyperosmotic stress, we focused on REMs, the best-characterized markers of plasma membrane nanodomains^42^ and known stabilizers of membrane topologies.^23^ Using live-cell laser scanning confocal microscopy in five-day-old seedling roots expressing YFP-REM1.2, we observed a diffuse yet polarized localization, with stronger fluorescence signals detected at the shootward regions of the plasma membrane (Figure 5A). Upon treatment with sorbitol for 5 minutes, YFP-REM1.2 formed nanodomains that are approximately 0.4 µm in diameter, and their density increased in a concentration-dependent manner with sorbitol (Figure 5A and 5B). This suggests that the density of YFP-REM1.2-positive nanodomains serves as a readout for the severity of hyperosmotic stress. Similar nanodomain formation was observed in other organs, including cotyledons and hypocotyls, when treated with sorbitol (Figure S5A). Because REMs are also components of plasmodesmata-localized nanodomains,^43^ we crossed a plasmodesmata marker, mCherry-PDCB1,^44^ with YFP-REM1.2 to study their co-association. At cell-cell junctions, a portion of YFP-REM1.2 nanodomains overlapped with mCherry-PDCB1 signal (Figure S5B), confirming their presence in plasmodesmata. However, YFP-REM1.2 nanodomains observed on the outer (periclinal) sides of cells were independent of plasmodesmata (Figure S5B). Our study focused on this latter population of YFP-REM1.2 nanodomains. Time-lapse imaging revealed rapid formation of these immobile nanodomains, likely instantaneously upon hyperosmotic stress (Figure S5C, Video S6). Notably, this process was reversible: the nanodomains dispersed back into the plasma membrane when hyperosmotic stress was alleviated (Video S6). Lastly, to evaluate the dependence of YFP-REM1.2 nanodomain formation on sterols, a key membrane lipid species that facilitates REM localization into highly ordered nanodomains^42^, we treated YFP-REM1.2 seedlings with fenpropimorph, a sterol biosynthesis inhibitor.^45^ Unlike the mock-treated control, fenpropimorph-treated seedlings did not exhibit YFP-REM1.2 nanodomains under hyperosmotic stress (Figure S5D). These results suggest that the formation of YFP-REM1.2 nanodomains relies on the sterol composition of the plasma membrane.

**Figure 5.**
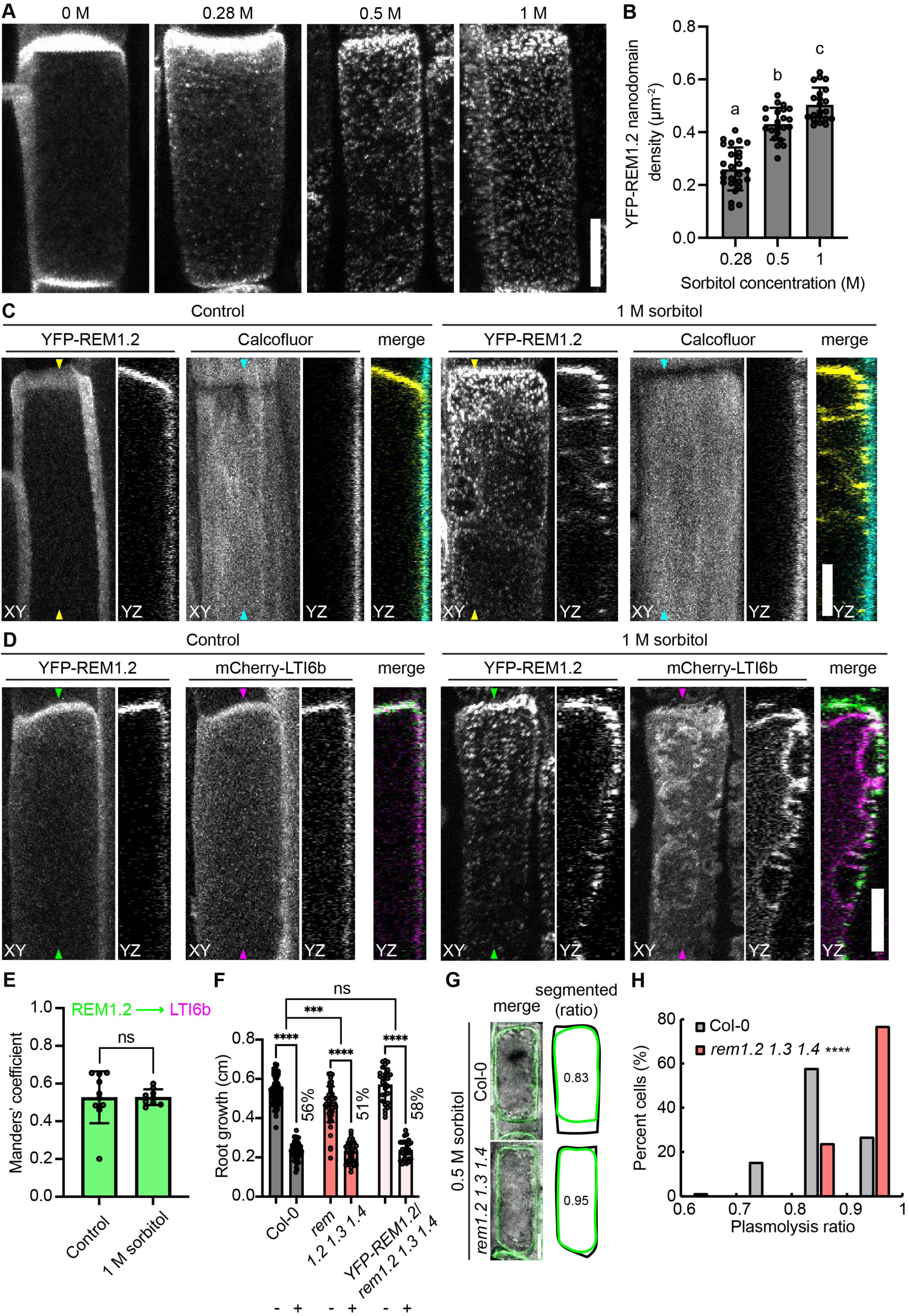
The rapidly forming REM nanodomains function as negative regulators of wall-membrane attachments. (A) Laser scanning confocal images in root epidermal cells of 5-d-old seedlings expressing YFP-REM1.2 treated with 0 M, 0.28 M, 0.5 M, or 1 M sorbitol for 5 min, respectively. Scale bar: 10 µm. (B) Bar graph showing YFP-REM1.2 nanodomain density at different sorbitol concentrations quantified from (A). *n* ≥ 21 cells from at least 5 seedlings per treatment. Error bars indicate SD. Different letters indicate significant differences by one-way ANOVA and Tukey’s test. (C) Laser scanning confocal images of root epidermal cells in 5-d-old seedlings expressing YFP-REM1.2 co-stained with calcofluor white under the control condition or treated with 1 M sorbitol for 5 min. XY and YZ projections are shown. The positions where YZ projections were made are indicated by arrowheads. Scale bar: 10 µm. (D) Laser scanning confocal images of root epidermal cells in 5-d-old seedlings expressing YFP-REM1.2 and mCherry-LTI6b under the control condition or treated with 1 M sorbitol for 5 min. XY and YZ projections are shown. The positions where YZ projections were made are indicated by arrowheads. Scale bar: 10 µm. (E) Quantification of Manders overlap coeffiicient in 5-d-old seedlings of YFP-REM1.2/mCherry-LTI6b under the control condition or treated with 1 M sorbitol for 5 min. Error bars indicate SD. *n* ≥ 9 cells per genotype per treatment. ns, no significance by Student’s *t*-test. (F) Root growth of wild type, *rem1.2 1.3 1.4* triple mutants, and *YFP-REM1.2*/*rem1.2 1.3 1.4* lines 1 day after transfer to either fresh MS media (-) or MS media supplemented with 0.28 M sorbitol (+). *n* ≥ 27 seedlings per genotype per treatment. Error bars indicate SD. ns, no significance; \*\*\**p* < 0.001, \*\*\*\**p* < 0.0001 by two-way ANOVA. (G) Laser scanning confocal images and manual tracings of plasmolyzed root epidermal cells in 5-d-old seedlings expressing YFP-LTI6b in Col-0 or *rem1.2 1.3 1.4* background, respectively. Plasmolysis was induced by 0.5 M sorbitol for 5 min. Each displayed ratio corresponds to the same individual cell. Scale bar: 10 µm. (H) Histogram showing plasmolysis ratio in root epidermal cells of 5-d-old seedlings expressing YFP-LTI6b in Col-0 or *rem1.2 1.3 1.4* background, respectively. *n* ≥ 102 cells from at least 10 seedlings per genotype. \*\*\*\**p* < 0.0001 by Student’s *t*-test.

To test whether YFP-REM1.2 nanodomains are associated with wall-membrane attachment sites, we performed live-cell imaging experiments using two approaches: staining five-day-old YFP-REM1.2 seedlings with calcofluor white (Figure 5C) and generating a YFP-REM1.2/mCherry-LTI6b double marker line (Figure 5D). Hyperosmotic stress-induced YFP-REM1.2 nanodomains were attached to the wall (Figure 5C) and remained colocalized with mCherry-LTI6b-labeled wall-membrane attachment sites at the outer surface of cells (Figure 5D and 5E). To investigate the role of REMs in hyperosmotic stress responses, we analyzed the *rem1.2*, *rem1.3*, *rem1.4* single mutants,^46^ the *rem1.2 1.3* double mutant,^43,46^ and the *rem1.2 1.3 1.4* triple mutant^46^ by assessing root growth and plasmolysis. The *rem* single mutants and the *rem1.2 1.3* double mutant did not differ from the wild type (Figure S5E-S5G), while *rem1.2 1.3 1.4* triple mutants displayed significantly reduced growth inhibition and resistance to plasmolysis under stress conditions (Figure 5F-5H), phenotypes that were similar to those of the *rhm1* and *shou4-3 4l-1* mutants (Figure 2D, Figure 3A and 3B, Figure 4G and 4H). Expression of YFP-REM1.2 under its native promoter in the *rem1.2 1.3 1.4* triple mutant background restored the root growth phenotypes to wild-type levels (Figure 5F), supporting functional redundancy among REM isoforms. Together, these results suggest that REMs rapidly form nanodomains upon hyperosmotic stress and REMs function to negatively affect wall-membrane attachments, though these two processes might be unrelated.

### REMs antagonize CSCs to regulate hyperosmotic stress responses

Given the observed formation of CSC clusters and REM nanodomains in root cells under hyperosmotic stress, we next sought to investigate their relationship. To visualize them, we crossed YFP-REM1.2 with tdTomato-CESA6 lines and imaged root cells using spinning disk confocal microscopy under control conditions or in the presence of 1 M sorbitol. Under control conditions, tdTomato-CESA6-labeled CSC particles were distributed across the plasma membrane, while YFP-REM1.2 signals were predominantly localized to the shootward regions of the plasma membrane (Figure 6A). Upon treatment with 1 M sorbitol, both markers formed distinct, punctate structures (Figure 6A and 6B). To quantify the spatial relationship of CSCs and REMs, we performed Manders’ colocalization analysis using the GFP-CESA3/tdTomato-CESA6 double marker line as a positive control (Figure S6A). Under control conditions, YFP-REM1.2 signals showed minimal overlap with tdTomato-CESA6-labeled CSC particles (Figure 6C). However, under hyperosmotic stress, REM1.2 nanodomains and CSC clusters exhibited partial colocalization, yet their Manders’ overlap coefficients remained significantly lower than those of the GFP-CESA3/tdTomato-CESA6 positive control (Figure 6C). These data suggest that CSCs and REMs function in spatially distinct cellular domains under hyperosmotic stress, though partial colocalization suggests there may be specific cellular contexts where their functions may be coordinated.

**Figure 6.**
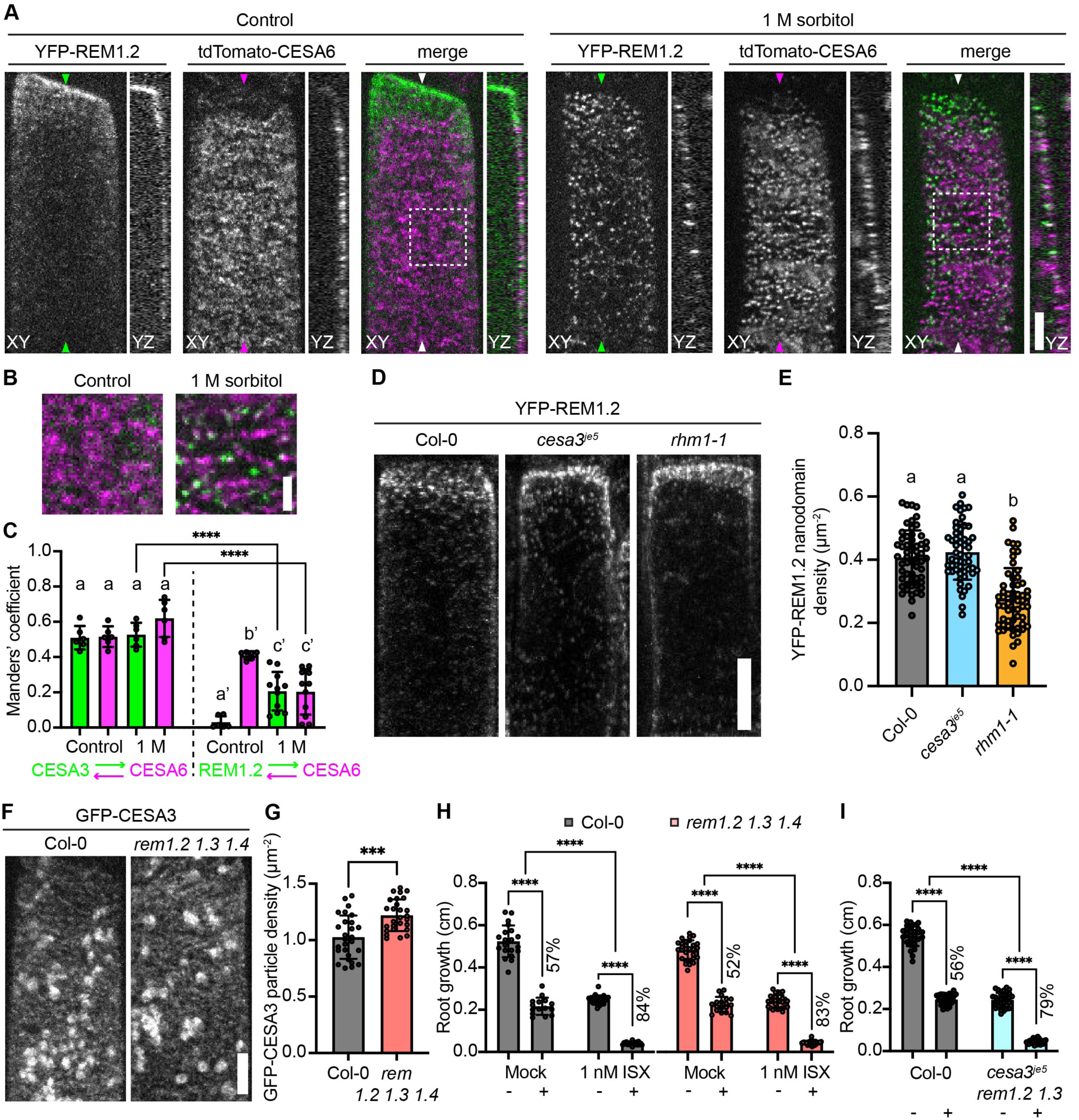
REMs antagonize CSCs to regulate hyperosmotic stress responses. (A) Spinning disk confocal images of root epidermal cells in 5-d-old seedlings expressing YFP-REM1.2 and tdTomato-CESA6 under the control condition or treated with 1 M sorbitol for 5 min. XY and YZ projections are shown. The positions where YZ projections were made are indicated by arrowheads. Scale bar: 5 µm. (B) Magnified views of the regions outlined by white dashed boxes in (A). Scale bar: 2 µm. (C) Quantification of Manders overlap coeffiicient in 5-d-old seedlings of GFP-CESA3/tdTomato-CESA6 or YFP-REM1.2/tdTomato-CESA6 under the control condition or treated with 1 M sorbitol for 5 min. Error bars indicate SD. *n* ≥ 6 cells per genotype per treatment. In each genotype, different letters indicate significant differences by one-way ANOVA and Tukey’s test. \*\*\*\**p* < 0.0001 by Student’s *t*-test when comparing the two genotypes under the same condition. (D) Laser scanning confocal images of YFP-REM1.2 nanodomains in 5-d-old seedlings of the Col, *cesa3^je^*^5^, or *rhm1-1* background, respectively when treated with 0.5 M sorbitol for 5 min. Scale bar: 10 µm. (E) Quantification of YFP-REM1.2 nanodomain density from (C). *n* ≥ 54 cells from at least 10 seedlings per genotype. Error bars indicate SD. Different letters indicate significant differences by one-way ANOVA and Tukey’s test. (F) Spinning disk confocal images of GFP-CESA3 in root epidermal cells of 5-d-old seedlings of Col and *rem1.2 1.3 1.4.* Scale bar: 5 µm. (G) Quantification of GFP-CESA3 particle density at the plasma membrane from (E). *n* = 27 cells from at least 6 seedlings per genotype. Error bars indicate SD. \*\*\**p* < 0.001 by Student’s *t*-test. (H) Effects of ISX on root growth in wild type and *rem1.2 1.3 1.4* mutants 1 day after transfer to either fresh MS media (-) or MS media supplemented with 0.28 M sorbitol (+) in the presence or absence of 1 nM ISX. *n* ≥ 15 seedlings per genotype per treatment. (I) Root growth of wild type and *cesa3^je^*^5^ *rem1.2 1.3* mutants 1 day after transfer to either fresh MS media (-) or MS media supplemented with 0.28 M sorbitol (+). *n* ≥ 28 seedlings per genotype per treatment. Error bars indicate SD. \*\*\*\**p* < 0.0001 by two-way ANOVA.

To determine whether there is a functional relationship between CSCs and REMs, we introduced the YFP-REM1.2 marker into the *cesa3^je^*^5^ and *rhm1-1* mutant backgrounds and measured REM nanodomain density following 0.5 M sorbitol treatment. While REM nanodomain density in the *cesa3^je^*^5^ mutant remained unchanged, it was significantly reduced in the *rhm1-1* mutant compared to the wild type (Figure 6D and 6E), suggesting that RHM1 promotes REM nanodomain formation. Additionally, we introduced the GFP-CESA3 marker into the *rem1.2 1.3 1.4* triple mutant and quantified CSC density at the plasma membrane under control conditions. *rem1.2 1.3 1.4* triple mutants exhibited significantly higher CSC density than wild type (Figure 6F and 6G). Given the importance of sterol biosynthesis for REM1.2 nanodomain formation (Figure S5D), we tested whether it also influences CSC patterning at the plasma membrane and affects plasmolysis. Compared to mock-treated roots, treatment with fenpropimorph had no detectable effect on CSC density at the plasma membrane or on protoplast behavior under 0.5 M sorbitol (Figure S6B-S6E). These data support the hypothesis that CSC density at the plasma membrane is necessary for plasma membrane-cell wall attachment, which plays a critical role in safeguarding root cells against the disruptive effect of hyperosmotic stress. Furthermore, our data suggest that REMs inhibit the formation of these attachments by limiting CSCs abundance at the plasma membrane, independently of REM nanodomain formation.

To test this hypothesis, we analyzed the effect of ISX on hyperosmotic stress-induced growth inhibition in *rem1.2 1.3 1.4* triple mutant roots. Under sorbitol treatment, mock-treated mutants exhibited a 52% reduction in growth, whereas ISX treatment exacerbated the reduction to 83% (Figure 6H), a level comparable to that observed in cellulose-deficient mutants (Figure 2A). We also generated the *cesa3^je^*^5^ *rem1.2 1.3* triple mutant and found that this line phenocopied the *cesa3^je^*^5^ single mutant under both control and hyperosmotic stress conditions (Figure 6I, Figure 2A). We further attempted to generate the *cesa3^je^*^5^ *rem1.2 1.3 1.4* quadruple mutant but were unsuccessful because the *cesa3^je^*^5^ and *rem1.4* mutations are located on the same arm of chromosome 5. Together, these results confirm that the reduced severity of hyperosmotic stress responses in *rem1.2 1.3 1.4* triple mutants is a consequence of enhanced CSC density at the plasma membrane, and further suggest an antagonistic relationship between REMs and CSCs.

### SHOU4/4L associate with REMs

To uncover additional proteins present in REM nanodomains that could mediate the apparent antagonism with CSCs, we employed proximity labeling and generated transgenic lines expressing *TbID-YFP-REM1.2* driven by the *REM1.2* native promoter in the wild-type background. After confirming that the TbID tag did not affect YFP-REM1.2 localization to the plasma membrane or nanodomain formation (Figure S6F), we treated five-day-old seedlings with 50 µM biotin for 15 minutes, in the presence or absence of 0.5 M sorbitol. In both conditions, biotinylated proteins were detected in addition to those that are endogenously biotinylated (Figure S6G and S6H). Overall, we identified 5,922 proteins, which were subjected to two filtering steps (Figure 7A). To eliminate background proteins, we compared TbID-YFP-REM1.2 samples with the wild-type control, identifying 30 and 111 proteins under the control and sorbitol-treated conditions, respectively (Figure 7B and 7C, Table S1). A subsequent comparison between these conditions revealed 83 proteins enriched specifically under sorbitol treatment (Figure 7D, Table S1).

**Figure 7.**
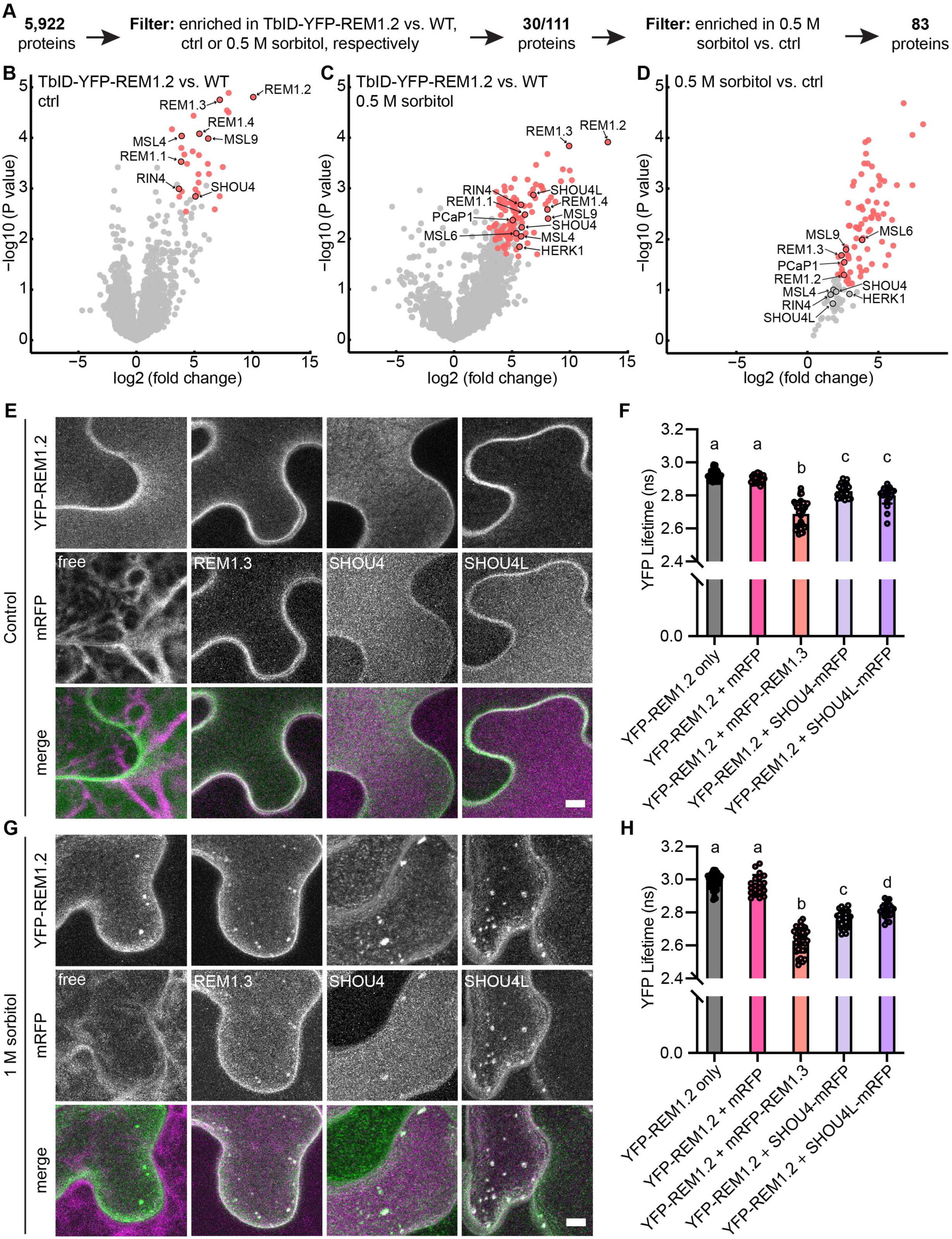
REM nanodomains harbor many other proteins in the vicinity, including SHOU4/4L. (A) Workflow of filtering steps to identify proteins in the vicinity of TbID-YFP-REM1.2 in the presence or absence of 0.5 M sorbitol. (B) and (C) Volcano plots showing differentially enriched proteins comparing TbID-YFP-REM1.2/Col-0 and Col-0 (WT) under the control (ctrl) condition (B) or treated with 0.5 M sorbitol for 15 min (C). Significantly enriched proteins are color coded using one-sided Welch’s *t*-test with an FDR of 0.05 and an S0 of 0.5. *n* ≥ 3 biological replicates. (D) Volcano plot showing differentially enriched proteins comparing 0.5 M sorbitol treatment and the control condition in TbID-YFP-REM1.2/Col-0. Pair-wise comparison was conducted using Student’s *t*-test with an FDR of 0.05 and an S0 of 0.5. *n* ≥ 3 biological replicates. (E) Laser scanning confocal images of tobacco leaf epidermal cells co-expressing YFP-REM1.2 with free mRFP, mRFP-REM1.3, SHOU4-mRFP, or SHOU4L-mRFP, respectively, under control conditions. Scale bar: 5 µm. (F) Measurements of YFP lifetime from (E). Error bars indicate SD. Different letters indicate significant differences by one-way ANOVA and Tukey’s test. *n* ≥ 23 regions of interest from at least 3 infiltrated plants per construct combination. (G) Laser scanning confocal images of tobacco leaf epidermal cells co-expressing YFP-REM1.2 with free mRFP, mRFP-REM1.3, SHOU4-mRFP, or SHOU4L-mRFP, respectively, upon 1 M sorbitol treatment for 10 min. Scale bar: 5 µm. (H) Measurements of YFP lifetime from (E). Error bars indicate SD. Different letters indicate significant differences by one-way ANOVA and Tukey’s test. *n* ≥ 20 regions of interest from at least 3 infiltrated plants per construct combination.

Our proteomic data revealed proteins known to interact with REM1.2, including REM1.2 itself, REM1.3 and other REMs in the same family,^46^ RPM1-INTERACTING PROTEIN4 (RIN4),^47^ and PLASMA-MEMBRANE ASSOCIATED CATION-BINDING PROTEIN1 (PCaP1)^48^ (Figure 7B-7D), confirming previous protein-protein interaction studies. Interestingly, some of these proteins did not meet the statistical threshold across all three pairwise comparisons. For instance, RIN4 was enriched in TbID-YFP-REM1.2 samples compared to the wild type under both control and sorbitol conditions, but not in the direct comparison between the two conditions (Figure 7B-7D). Similarly, PCaP1 was enriched in TbID-YFP-REM1.2 samples compared to the wild type under sorbitol conditions, but not under control conditions; however, it was enriched in the comparison between the two conditions (Figure 7B-7D). These findings raise the possibility of false negatives in the pairwise comparisons described above and led us to test whether these proteins colocalize and interact with REM1.2 during hyperosmotic stress using an orthogonal approach. To this end, we performed confocal microscopy and FRET-FLIM analyses by transiently expressing YFP-REM1.2 in tobacco leaves, either alone or alongside an mRFP-tagged protein of interest. mRFP-REM1.3 served as a positive control, while free mRFP was used as a negative control. As anticipated, free mRFP localized to the cytosol did not colocalize with or interact with YFP-REM1.2 under either control or sorbitol treatment conditions (Figure 7E-7H). mRFP-REM1.3 colocalized and interacted with YFP-REM1.2 at the plasma membrane under control conditions and within nanodomains following sorbitol treatment (Figure 7E-7H). Similarly, mRFP-RIN4 and PCaP1-mRFP also exhibited colocalization and interaction with YFP-REM1.2 under both control and sorbitol conditions (Figure S7A-S7D).

One group of proteins identified in the pairwise comparisons was the MECHANOSENSITIVE CHANNEL OF SMALL CONDUCTANCE-LIKEs (MSLs), including MSL4, MSL6, and MSL9 (Figure 7B-7D). In tobacco infiltration assays, mRFP-MSL4 and mRFP-MSL6 colocalized with YFP-REM1.2 at the plasma membrane under control conditions but not within nanodomains following sorbitol treatment (Figure S7E and S7F). However, under prolonged sorbitol treatment, colocalization was observed in the Hechtian reticulum (Figure S7E and S7F). In contrast, the localization pattern of mRFP-MSL9 resembled that of the cortical endoplasmic reticulum (ER) under both control and sorbitol conditions (Figure S7G), consistent with the close proximity of the ER to the plasma membrane where YFP-REM1.2 is located. This observation aligns with previous reports of MSL10, the closest homolog of MSL9 in the MSL family, being localized at ER-plasma membrane contact sites.^49,50^ To assess the functional role of MSLs in Arabidopsis root growth responses to hyperosmotic stress, we compared the growth phenotypes of the *msl4;5;6;9;10* quintuple mutant and the wild type. However, no significant differences in growth were observed between the two genotypes under either control or sorbitol treatment conditions (Figure S7H), suggesting that none of these MSLs are required for root growth responses to hyperosmotic stress.

Another group of proteins identified in the pairwise comparisons was SHOU4/4L, known inhibitors of CSC exocytosis.^26^ Similar to the positive control mRFP-REM1.3, both SHOU4-mRFP and SHOU4L-mRFP colocalized and interacted with YFP-REM1.2 at the plasma membrane under control conditions and within nanodomains following sorbitol treatment (Figure 7E-7H, Figure S7I). In Arabidopsis roots, the *shou4-3 4l-1* double mutants displayed reduced root growth inhibition and increased resistance to plasmolysis than wild type under hyperosmotic stress (Figure 4G and 4H). These phenotypes closely resembled those of the *rem1.2 1.3 1.4* triple mutants (Figure 5F-5H). Together, these findings suggest that SHOU4/4L associate with REMs to mediate the antagonism with CSCs.

## Discussion

In summary, we provided genetic, live-cell imaging, and proximity labeling-based proteomic evidence that CSCs and REMs mediate the association and function of the cell wall-plasma membrane interface. While they are prominently observed at the outer epidermal cell surface where they are in direct contact with the environment and function independently of plasmodesmata, they are also likely present and functionally relevant at other wall-membrane interfaces, such as in inner tissues where high-resolution live-cell imaging is technically challenging. Maintaining CSC density at the plasma membrane is crucial for proper cell wall-plasma membrane attachment and root responses to hyperosmotic stress. Baseline CSC density under normal conditions sets the capacity for CSC relocalization and clustering at wall-membrane attachment sites during hyperosmotic stress. This CSC-dependent reorganization promotes wall-membrane attachment that both limits membrane detachment during plasmolysis and enables the efficient resumption of growth following stress. In parallel, REMs rapidly form nanodomains at the plasma membrane upon hyperosmotic stress and they harbor many additional proteins in the vicinity, including the CSC exocytosis inhibitors SHOU4/4L.^26^ These findings reveal a molecular module in which CSCs are the core component that mediate attachment between the cell wall and plasma membrane, acting downstream of and being antagonized by REMs. Our findings provide new insights into the structural determinants of wall-membrane attachment in plants and a physiological context in which their function supports the resilience of plants to environmental stress.

Our survey of a panel of cell wall mutants in hyperosmotic stress-induced growth inhibition revealed how different classes of cell wall polysaccharides are involved in this process. This analysis led us to focus on two key proteins involved in cell wall biosynthesis: CSCs, and RHM1. RHM1 encodes a UDP-L-rhamnose synthase, initially identified as a suppressor of LEUCINE RICH REPEAT-EXTENSIN1 (LRX1), an extracellular protein that regulates cell wall formation in root hairs.^51,52^ RHM1 plays an important role in the morphogenesis of epidermal cells, including root hairs,^51^ cotyledon pavement cells,^53^ and conical petal cells.^28,54^ Although rhamnose is required for both pectin biosynthesis and flavonol glycosylation, experimental evidence suggests that rhamnose-containing RG-I directly contributes to epidermal morphogenesis, whereas flavonol rhamnosylation has an indirect effect.^53,55^ Our data reveal a previously uncharacterized function of RHM1 in regulating root growth and responses to hyperosmotic stress (Figures 2, 3, S1, and S2). While we cannot definitively determine whether the observed *rhm1* mutant phenotypes in our growth and plasmolysis assays result directly from RG-I rhamnose deficiency, we argue that an indirect mechanism is more likely. This is supported by our findings that CSC machinery components are enriched, and CSC density at the plasma membrane is increased, in the *rhm1-1* mutant (Figures 3, S2J, and S3F). Additionally, the *cesa3* mutation is epistatic to *rhm1-1* (Figure 4I). These findings underscore the intricate interplay among cell wall polymers:^7^ altering one wall component can influence the synthesis, assembly, and/or organization of others.^56–58^

Environmental changes influence CSC dynamics and cellulose synthesis.^59^ For example, salt stress reduces cellulose production and leads to CSC depletion and reappearance at the plasma membrane that coincides with cortical microtubule reorganization,^60^ a process requiring COMPANION OF CELLULOSE SYNTHASE proteins^61^ and TETRATRICOPEPTIDE THIOREDOXIN-LIKE (TTL) proteins.^62^ Treating Arabidopsis seedlings with hyperosmotic stress for several hours, whether mild (200 mM mannitol) or moderate (500 mM mannitol), triggers the redistribution of CSCs from the plasma membrane into small CESA compartments (SmaCCs)^40^ or microtubule-associated cellulose synthase compartments (MASCs).^63^ Prolonged severe osmotic stress has also been shown to reduce cellulose production.^64^ In alignment with these previous observations, we demonstrate that acute, severe osmotic stress causes a subset of CSCs to remain at the plasma membrane, decorating Hechtian structures at cell wall-plasma membrane attachment sites (Figures 4 and S3, Videos S4 and S5). These CSCs are immobile (Figure S3, Video S3) and are likely inactive in cellulose synthesis, as CSC movement at the plasma membrane is thought to be driven by cellulose polymerization.^18^ Importantly, Alexa568-tdCBM3a labeling reveals that these plasma membrane-retained CSCs remain associated with cellulose at the cell surface under both control and hyperosmotic conditions (Figure S3A), which would be most consistent with the hypothesis that the CSC particles remain attached to the wall through their nascently synthesized cellulose.^19^ Importantly, we find that CSC density at the plasma membrane under non-stress conditions establishes a baseline that predicts CSC retention under hyperosmotic stress (Figure 3F-3H, Figure S3, Figure S4J and S4K, Video S5), linking steady-state CSC distribution to stress-induced reorganization outcomes. Consistent with this idea, we further demonstrate that CSC density correlates with both the extent of root cell plasmolysis and organ-level growth responses, including the capacity for growth recovery following stress removal (Figures 2, 3, 4, 5, S1, S2, and S4). In addition, wall-membrane attachment sites visualized by both light microscopy and cryogenic electron tomography show that the distal ends of Hechtian structures are closely associated with cellulose microfibrils within the cell wall (Figures 1 and S1A, Videos S1 and S2). Together, these findings provide direct experimental evidence for the longstanding hypothesis that CSCs contribute to plasma membrane anchoring to the cell wall, likely via nascent cellulose microfibrils integrated into the wall.^19^

REMs, a plant-specific protein family,^65^ are known to function in development,^43,46^ plasmodesmata aperture regulation,^43^ membrane topology stabilization in root nodule symbiosis,^23^ and plant innate immunity.^66^ Our work expands the understanding of REM function in environmental stress responses by demonstrating that the sterol-dependent (Figure S5), rapid formation of REM nanodomains at wall-membrane attachment sites (Figure 5, Video S6) upon hyperosmotic stress serves as a hub for the aggregation of many other proteins (Figures 7 and S7). This is in line with a recent preprint showing that salinity similarly induces REM1.2 nanodomain formation.^67^ In addition, we show that REMs act upstream of CSCs, with SHOU4/4L as their mediator, to regulate hyperosmotic stress responses in elongating roots (Figures 6 and 7). The elevated CSC density observed in *rem1.2 1.3 1.4* triple mutants (Figure 6) and in *shou4-3 4l-1* double mutants (Figure S4J and S4K),^26^ together with enhanced growth recovery following stress in *shou4-3 4l-1* double mutants (Figure S4L), indicates that increased baseline CSC density confers greater resistance to CSC depletion and promotes post-stress growth restoration. The unchanged CSC density following fenpropimorph treatment (Figure S6), along with the interaction between SHOU4/4L and REM1.2 in the absence of stress (Figures 7 and S7), suggests that REM-CSC antagonism occurs independently of REM nanodomain formation. This plasma membrane-localized molecular module, comprising CSCs, SHOU4/4L, and REMs, functions both as a structural anchor linking the cell wall to the plasma membrane and as a regulatory platform for coordinating hyperosmotic stress responses.

### Limitations of the Study

The ultrastructural and organizational details of how CSCs mediate the physical association between the cell wall and plasma membrane remain to be fully resolved. Although REM proteins were initially characterized for their ability to bind both simple and complex galacturonides *in vitro*,^24^ it is still unclear whether such interactions occur *in vivo*. If they do, how REMs, peripheral proteins anchored to the inner leaflet of the plasma membrane, engage with cell wall pectins remains an open question. Our data reveal that CSC density at the plasma membrane under non-stress conditions establishes a baseline that predicts CSC retention during hyperosmotic stress, and that this baseline correlates with both the extent of plasmolysis at the cellular level and growth inhibition and recovery at the organ level. While these findings suggest a functional link between wall-membrane attachment, plasmolysis, and growth resilience following stress, the causal relationships among these processes remain to be understood. In addition, proximity labeling identified numerous proteins in the vicinity of REM1.2, yet how these interactions change in response to hyperosmotic stress remains to be explored. Finally, the mechanical properties of both the cell wall and the plasma membrane, and how they are modulated by CSC density and REM-dependent organization, remain to be explored and represent an important dimension for understanding plant responses to the mechanical stress of a hyperosmotic environment. These questions present exciting future directions that we aim to pursue through interdisciplinary approaches to advance our understanding of plant-environment interactions.

## Resource Availability

### Lead contact

Further information and requests for resources and reagents should be directed to and will be fulfilled by the lead contact, José Dinneny.

### Material availability

Constructs and plant seeds generated in this study will be available from the lead contact upon request.

### Data and code availability

All data required to substantiate the claims of this paper are included in main or supplemental data. The full proteomics datasets are available in the PRIDE database (https://www.ebi.ac.uk/pride) under the dataset identifiers PXD065856, PXD065860, and PXD065862. This paper does not report original code. Any additional information required to reanalyze the data reported in this paper is available from the lead contact upon request.

## Acknowledgements

We thank past and present members of the Dinneny lab, the Plant Biology community at Stanford University, Charles T. Anderson and his group at The Pennsylvania State University, and the glycoscience research community at the Complex Carbohydrate Research Center at the University of Georgia for insightful discussions and critical feedback. We are especially grateful to Andrea Mair and Dominique Bergmann for generously providing proximity labeling constructs and offering valuable advice on proximity labeling experiments. We thank Renate Weizbauer, Andrey Malkovskiy, and the Carnegie Advanced Imaging Facility for access to and assistance with the spinning disk confocal microscope. We also thank Daniel Miranda and Olivia Creasey for their help with 3D segmentation using AIVIA software, and Yijia Yang for assistance with Arabidopsis genotyping. We gratefully acknowledge the following colleagues for sharing genetic materials: Siamsa Doyle and Stéphanie Robert for *mur2-1*, *mur3-1,* and *mur4-1* seeds; Olivier Hamant for *qua2-1* seeds; Henrik Scheller for *tbl10-1* and *gals1/2/3* seeds; Christoph Ringli for *RHM1-GFP*/*rhm1-2* seeds; Adam Saffer and Vivian Irish for *rhm1-3* seeds; Renee Weizbauer and David Ehrhardt for GFP-CESA3/mCherry-TUA5 seeds; Yvon Jaillais for mCherry-LTI6b seeds; Yoselin Benitez-Alfonso for mCherry-PDCB1 seeds; Xu Chen for *rem1.2 1.3c* seeds; and Charles T. Anderson for GFP-CESA3/tdTomato-CESA6 seeds.

Y.R. is a 2020 Simons fellow of the Life Sciences Research Foundation. This project was supported by grants awarded to J.R.D. from the Biological and Environmental Research (BER) Program, US Department of Energy (DE-AC02-76SF00515), National Institutes of Health, National Institute of General Medicine Sciences (R01 GM123259-01), the Stanford Precourt Institute, and a Faculty Scholars grant from the Simons Foundation and Howard Hughes Medical Institute (55108515). J.R.D. is an investigator of the Howard Hughes Medical Institute. M.Z. and P.D.D. were supported by the US Department of Energy, Office of Science, Office of Biological and Environmental Research under Award Number DE-AC02-76SF00515 (FWP 100883). N.B.A and T.O. were supported by the German Research Foundation (Deutsche Forschungsgemeinschaft; DFG) in frame of the Collaborative Research Center 924 (project ID 170483403). S.P.S.C. was supported by US Department of Energy award DE-SC0019313 and US National Science Foundation Division of Chemical, Bioengineering, Environmental and Transport Systems (CBET) Career Award (1846797). J.J.K. was supported by the National Science Foundation (IOS-1856431). S.-L.X. was supported by the National Institutes of Health (S10OD030441) and the Carnegie endowment fund to the Carnegie mass spectrometry facility.

## Author Contributions

Conceptualization, Y.R. and J.R.D.; data acquisition and analysis, Y.R., M.Z., W.P.D., A.V.R., T.S.G., N.B.A, D.J., and S.-L.X.; data interpretation, all authors; funding, Y.R., S.P.S.C., T.O., J.J.K., P.D.D., S.-L.X., and J.R.D; supervision, Y.R. and J.R.D; writing of the original draft, Y.R. and J.R.D; manuscript editing, Y.R., M.Z., W.P.D., N.B.A, D.J., S.P.S.C., T.O., J.J.K., P.D.D., S.-L.X., and J.R.D..

## STAR Methods

### Key resources table

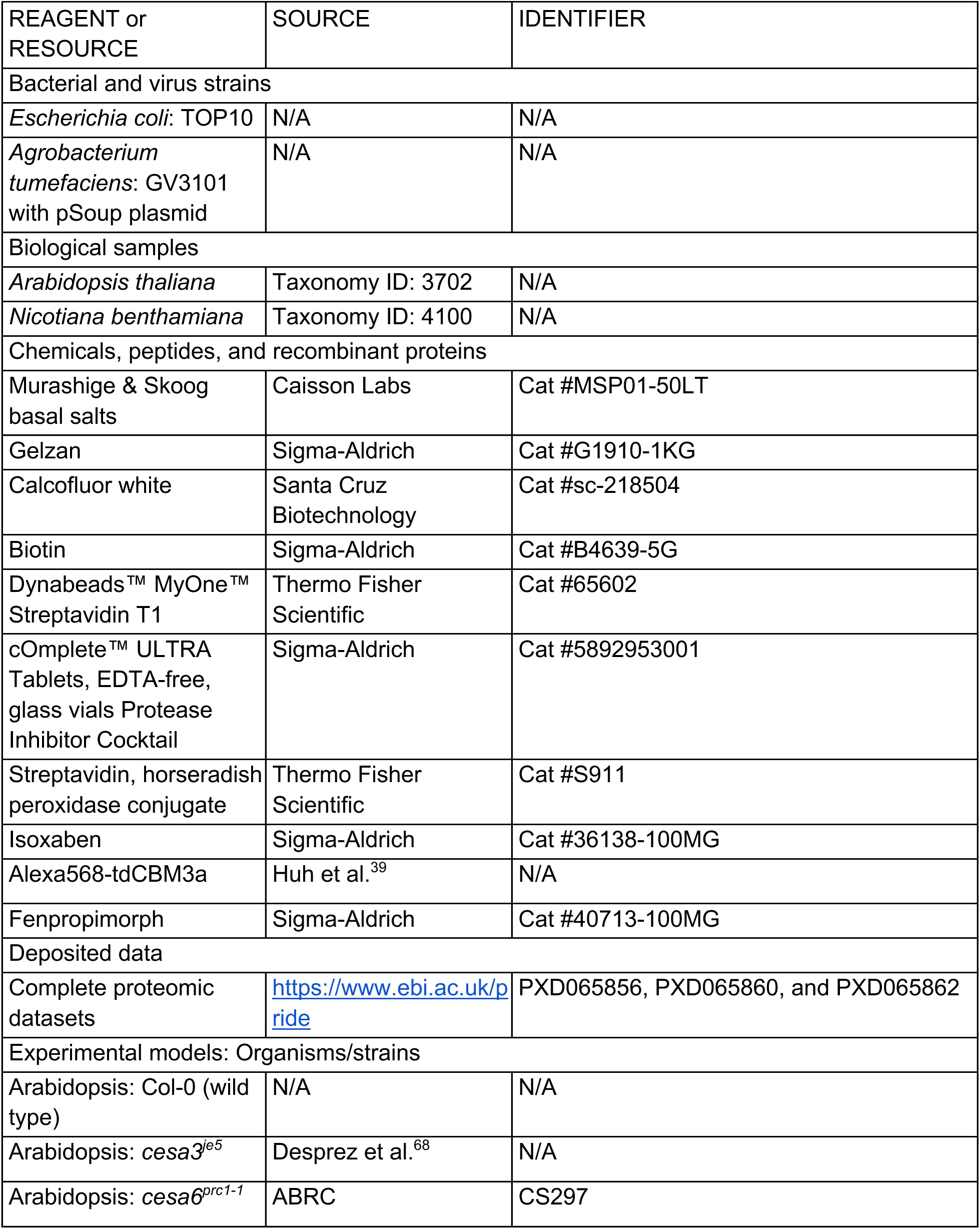
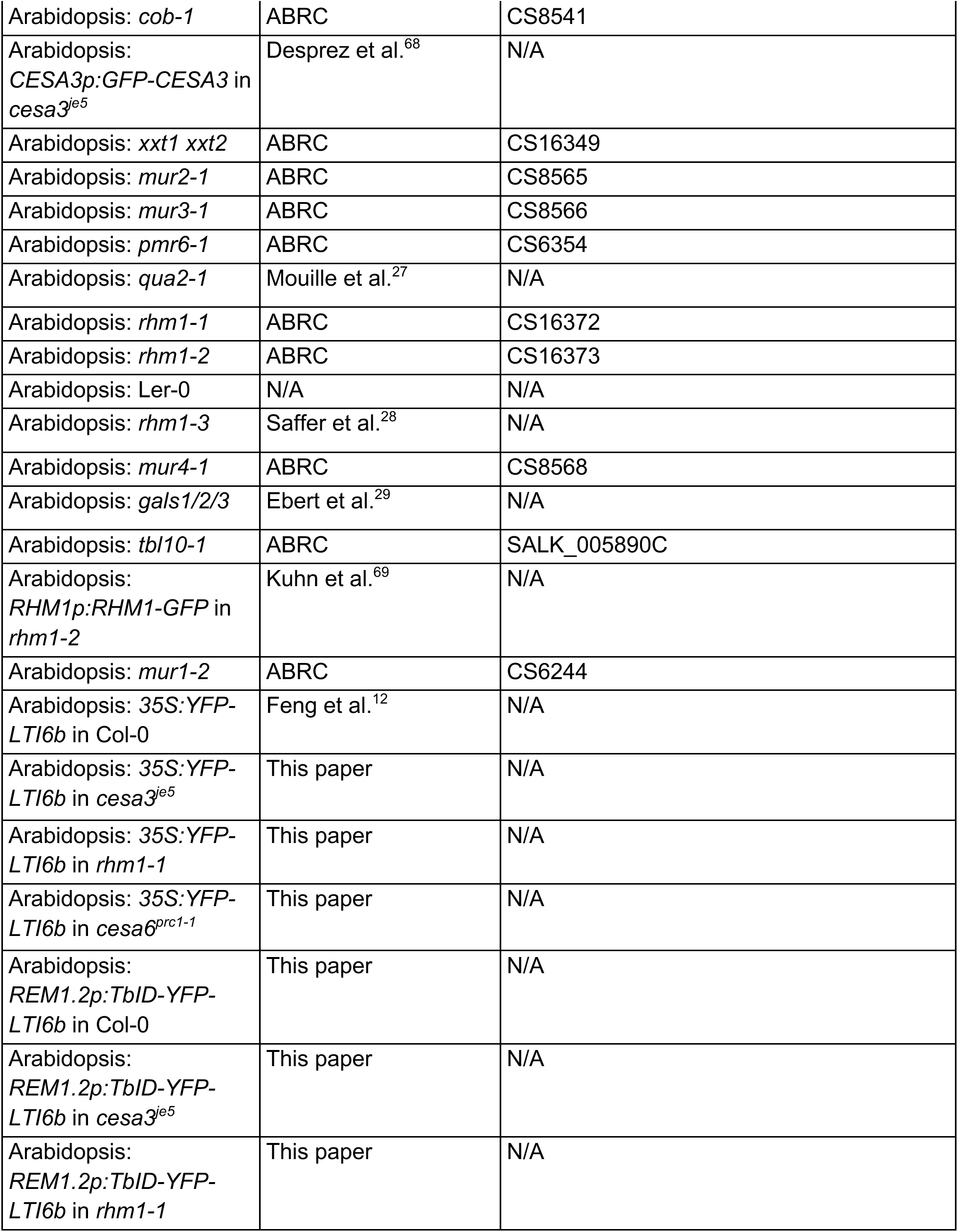
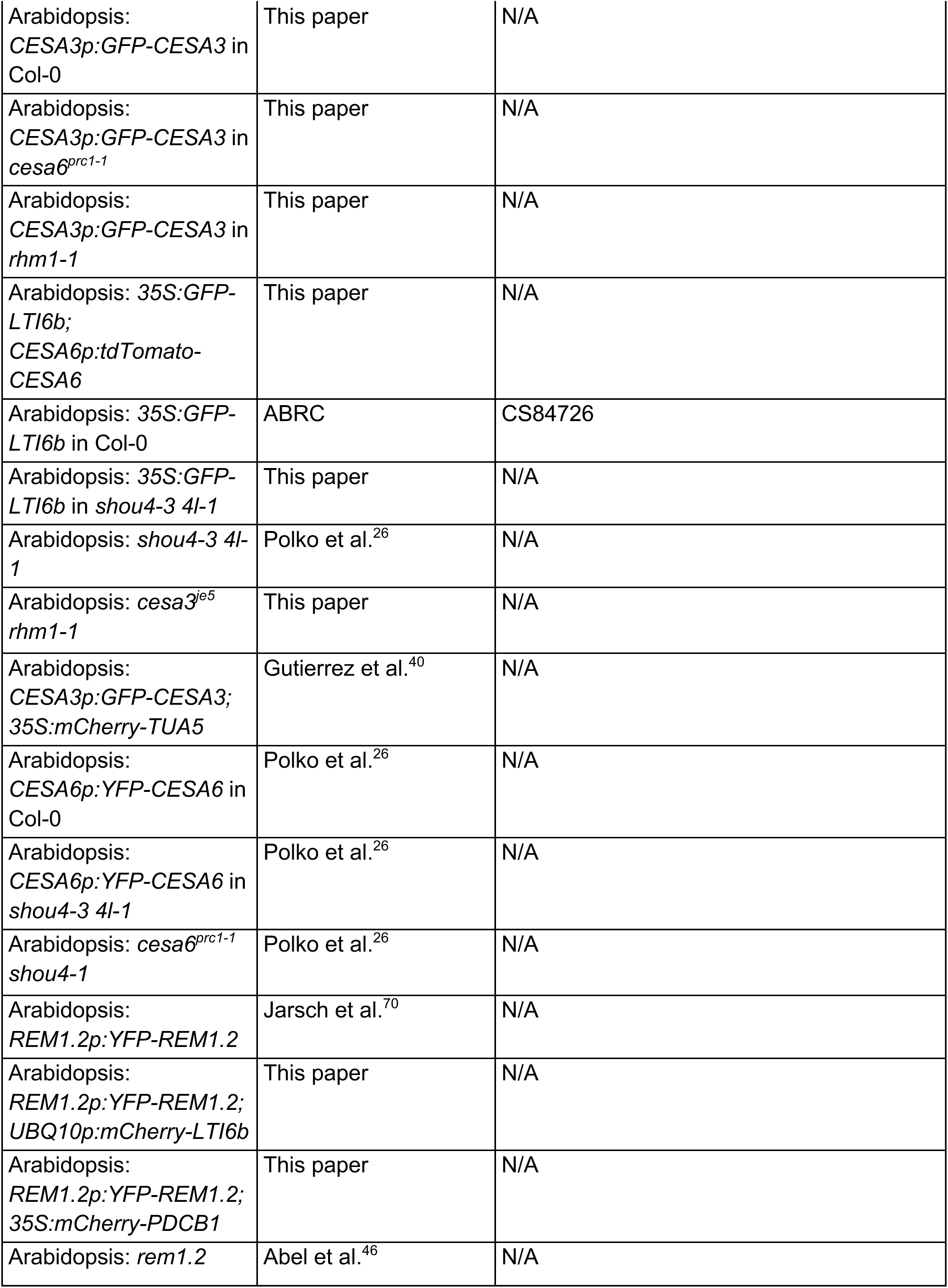
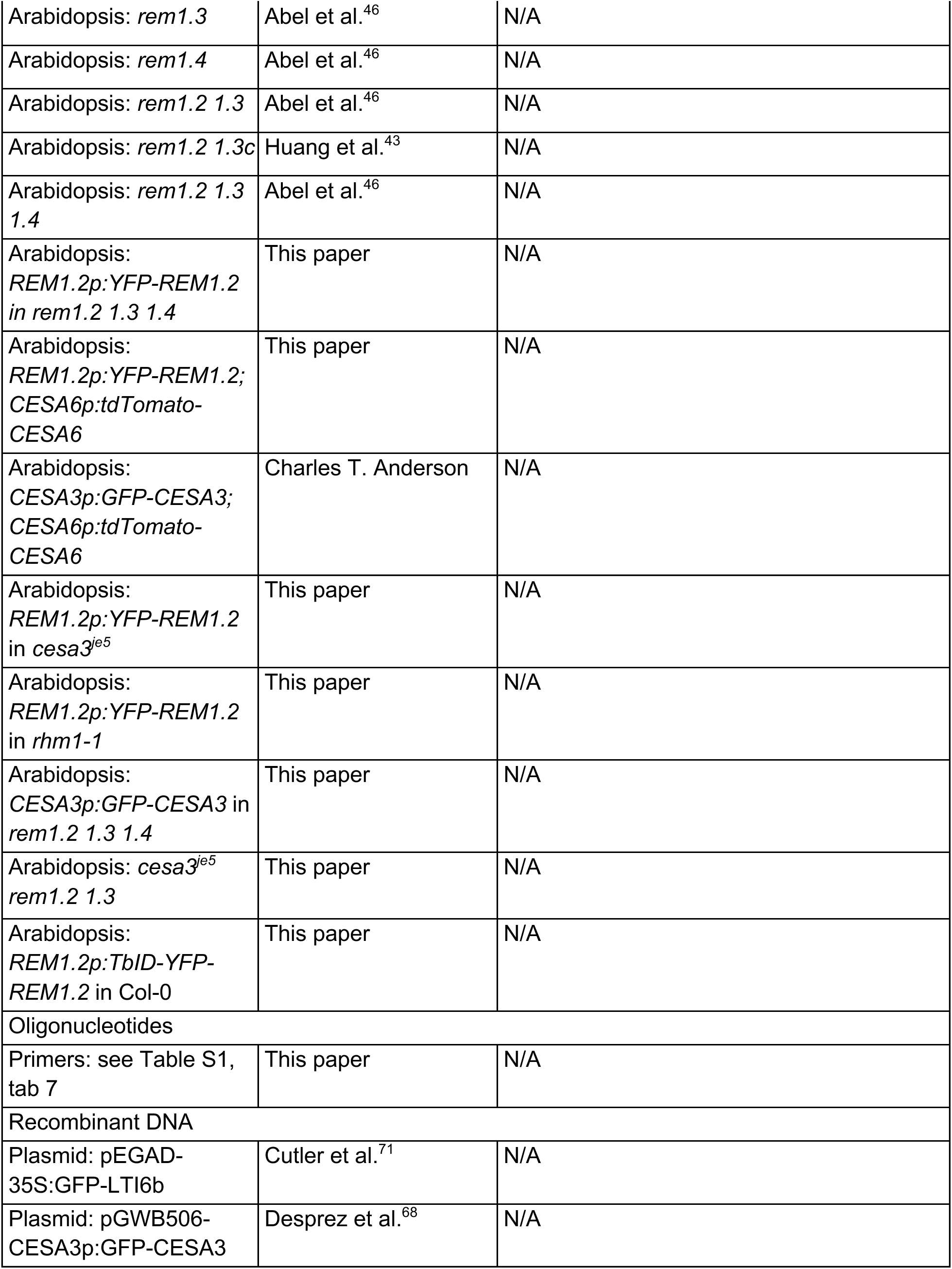
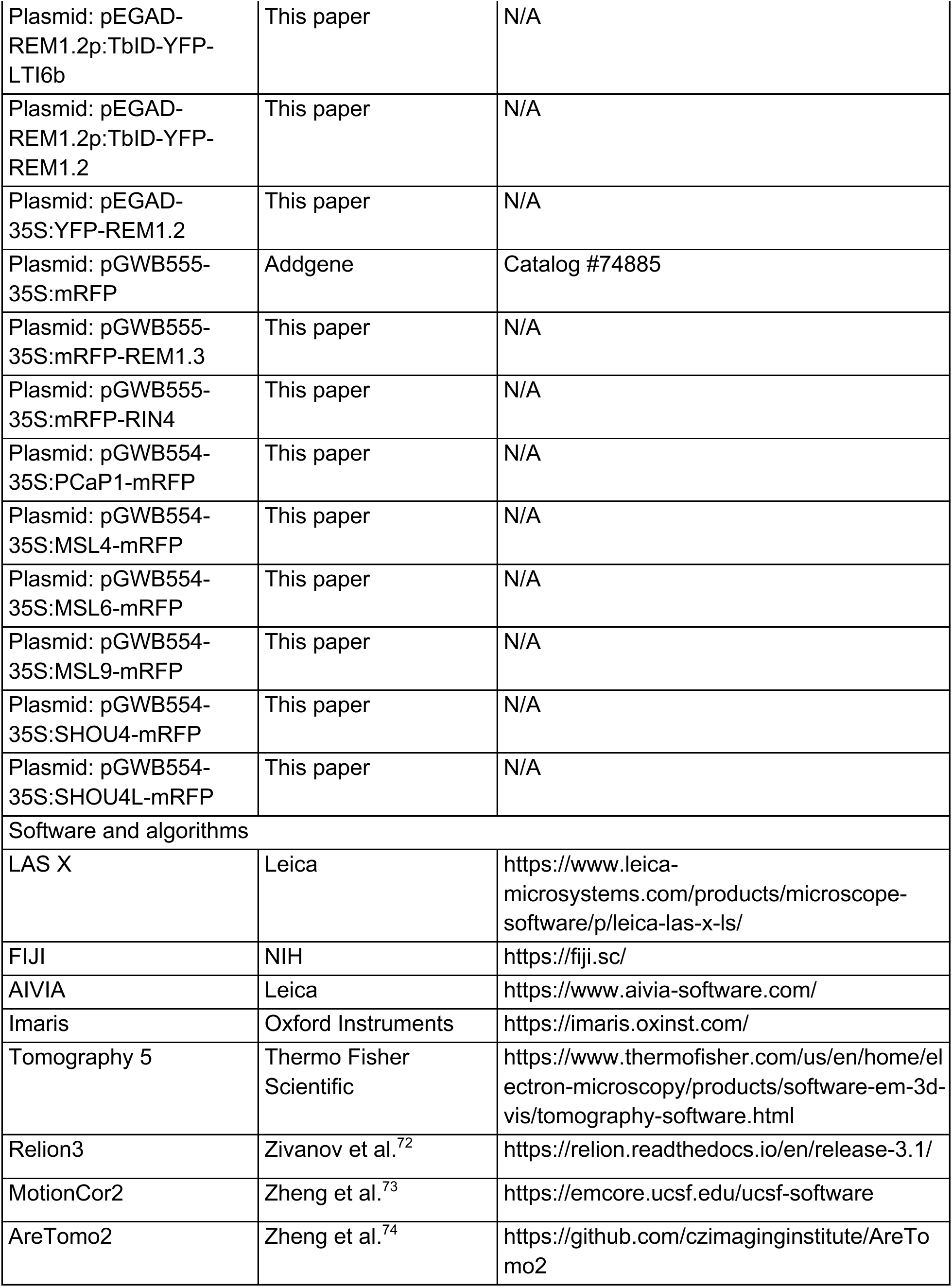
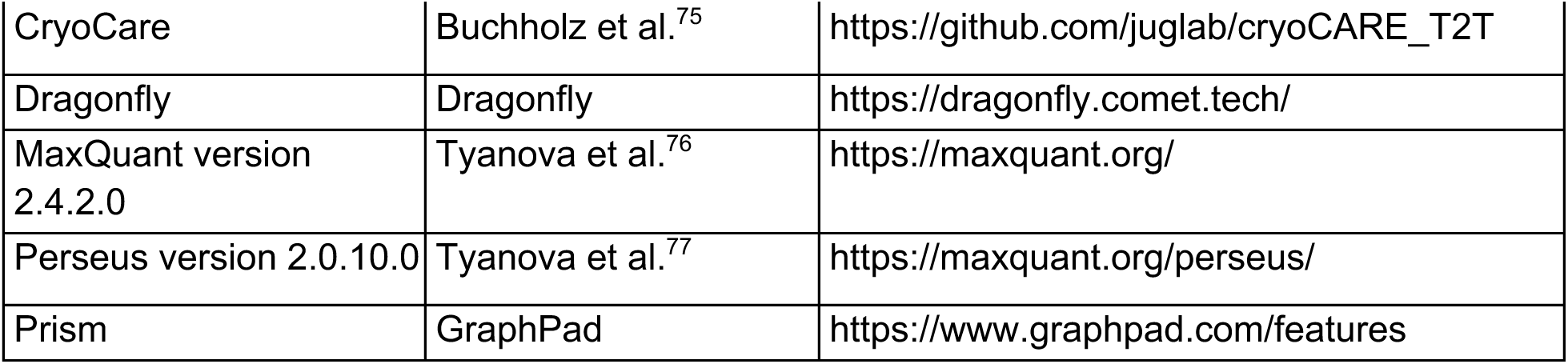

Experimental model and study participant details

- *Arabidopsis thaliana*, seedlings. Refer to the Key Resources Table for details on the lines used, and to the Method details section for growth conditions and experimental procedures.
- *Nicotiana benthamiana*, leaves. Refer to the Method details section for growth conditions and experimental procedures.

## Method details

### Plant Materials and Growth

*Arabidopsis thaliana* seeds were surface-sterilized using a solution of 30% bleach and 0.1% SDS for 20 minutes, followed by four rinses with sterile water. The sterilized seeds were stratified in the dark at 4°C for 2-3 days before being germinated on 1X Murashige and Skoog (MS) plates supplemented with 1% sucrose and solidified with 0.7% gelzan (pH 5.7). Seedlings were grown vertically in a growth chamber maintained at 22°C under a 16-hour light/8-hour dark photoperiod.

### Molecular Cloning and Plant Transformation

The *CESA3* coding sequence with attB sites was amplified from Arabidopsis Col-0 cDNA and introduced into the pDONR221 vector (Thermo Fisher, catalog #12536017) using a Gateway BP reaction. A 1831-bp promoter fragment upstream of the *CESA3* start codon was amplified from Col-0 genomic DNA with SbfI and XbaI restriction sites. The destination vector pGWB506 (Addgene, catalog #74848) was digested with SbfI and XbaI to replace the 35S promoter with the *CESA3* promoter. The *CESA3* coding sequence was subsequently inserted into the modified pGWB506 vector via a Gateway LR reaction to generate the *ProCESA3:GFP-CESA3* construct. This construct was transformed into Arabidopsis Col-0 plants, and transgenic lines were selected on medium containing 25 µg/mL Hygromycin B.

TurboID and YFP were amplified from the plasmid pENTR_L1-YFP-Turbo-L2 (Addgene, catalog #127350). *LTI6b* was amplified from Col-0 genomic DNA, including its exons, introns, and 3’UTR. *REM1.2* was amplified from Col-0 cDNA. To generate TbID-YFP-LTI6b and TbID-YFP-REM1.2, overlapping PCR was performed using these amplified fragments, introducing AgeI and EcoRI restriction sites at the N- and C-termini, respectively. A 1568-bp promoter fragment upstream of the *REM1.2* start codon was amplified from Col-0 genomic DNA, incorporating PmeI and AgeI sites. The pEGAD empty vector (ABRC stock #CD3-389) was digested with PmeI and AgeI to replace the *35S* promoter with the *REM1.2* promoter. This modified pEGAD vector was subsequently digested with AgeI and EcoRI to replace the GFP sequence with TbID-YFP-LTI6b or TbID-YFP-REM1.2. Constructs were then transformed into Arabidopsis Col-0 or mutant backgrounds to generate transgenic lines, which were then selected using 50 µM Basta.

The *35S:YFP-REM1.2* construct used in tobacco infiltration assays was generated following the same restriction digestion strategy described above. For *35S* promoter-driven mRFP-tagged protein constructs, the coding sequences of *REM1.3*, *RIN4*, *PCaP1*, *SHOU4*, and *SHOU4L* were amplified from Col-0 cDNA with attB sites. The genomic sequences of *MSL4*, *MSL6*, and *MSL9*, including exons and introns, were amplified from Col-0 genomic DNA with attB sites. All amplified sequences were first cloned into the pDONR221 vector and subsequently introduced into pGWB554 (Addgene, catalog #74884) or pGWB555 (Addgene, catalog #74885) to create mRFP fusion constructs with C- or N-terminal tags, respectively.

### Root Growth Assay

Four-day-old seedlings grown on standard 1X MS media were transferred to either fresh standard media or media supplemented with 0.28 M sorbitol. At the time of transfer, root tips were marked to track growth. Seedlings were then grown for an additional day, after which the plates were scanned using a dual-light flatbed scanner (Epson V800). Root growth was quantified in ImageJ by measuring the root length extending beyond the marked position. For growth rate analysis (Figure S1C and S1D), root growth was recorded using a custom live-imaging system as described previously.^78^ Growth rates were then measured manually in ImageJ.

For the organ-level growth recovery assay, four-day-old seedlings grown on standard 1X MS media were transferred to media supplemented with 0.5 M sorbitol for 30 min to induce an acute osmotic shock, or to standard media as a control. Seedlings were then transferred back to fresh standard media and allowed to grow for an additional day. Root growth was quantified as described above by measuring the root length extending beyond the marked position in scanned images.

### Pharmacological Treatments

To inhibit cellulose biosynthesis, we used isoxaben (ISX) (Sigma, catalog #36138-100MG) in two different experimental setups. For acute clearance of CSCs from the plasma membrane, five-day-old seedlings expressing GFP-LTI6b and tdTomato-CESA6 were incubated in MS liquid containing 100 nM ISX for 20 min prior to imaging for plasmolysis. MS liquid with 0.1% DMSO served as the mock treatment. To assess the effect of ISX on root growth, four-day-old seedlings were transferred from standard 1X MS plates to one of four treatment conditions: standard plates with 0.1% DMSO, standard plates with 1 nM ISX, 0.28 M sorbitol plates with 0.1% DMSO, or 0.28 M sorbitol plates with 1 nM ISX. Root growth was subsequently measured as described above. To analyze the effect of ISX on root growth recovery, four-day-old seedlings were transferred to media supplemented with 0.5 M sorbitol for 30 min to induce an acute osmotic shock, or to standard media as a control. Seedlings were then transferred back to one of two treatment conditions: standard plates with 0.1% DMSO and standard plates with 1 nM ISX. Root growth was then measured as described above.

To inhibit sterol biosynthesis, 100 µg/mL fenpropimorph (Sigma, catalog #40713-100MG) was added to standard 1X MS media. 1X MS media containing 0.1% DMSO was used as the mock. Four-day-old seedlings expressing YFP-REM1.2 were transferred from standard plate to either the mock or fenpropimorph plate and let grow for an additional day.

### Confocal Microscopy

To visualize plasmolysis in root cells in transgenic lines expressing YFP- or GFP-LTI6b, five-day-old seedlings were treated with 0.5 M sorbitol for 5 min on a glass slide and imaged on a Leica TCS SP8 microscope with an HC PL APO 93x/1.3 NA objective. YFP was excited with a white light laser at 514 nm, with emission detected between 525-570 nm. GFP was excited at 488 nm, with emission detected between 500-560 nm. Bright-field and fluorescence images were acquired simultaneously. The plasmolysis ratio was quantified in ImageJ by dividing the protoplast area (measured from the fluorescence image) by the total cell area (measured from the bright-field image). Protoplast curvature at cell ends was measured following previously established methods.^79^ Plasmolysis in etiolated hypocotyls was performed by treating three-day-old seedlings with 1 M sorbitol for 1 h on a glass slide and imaged as described above.

For growth recovery at the cellular level, five-day-old seedlings expressing YFP-LTI6b were treated with 0.5 M sorbitol for 5 min on a glass slide. Seedlings were then transferred to a gel pad prepared from 1X MS medium, sandwiched between two coverslips, and imaged immediately using a Leica TCS SP8 microscope with an HC PL APO 40x/1.10 NA objective. Following imaging, seedlings were returned to the growth chamber and allowed to grow vertically for 1 h, after which they were re-imaged using the same settings. Cell elongation was quantified as the change in length of the same cells between the two time points.

For calcofluor white staining, five-day-old seedlings were stained with 0.1 µM (10^-5^%, w/v) calcofluor white (Santa Cruz Biotechnology, catalog #sc-218504) diluted in MS liquid in the dark for 30 min, washed with MS liquid, treated with fresh MS liquid or 1 M sorbitol for 5 min on a glass slide, and imaged on a Leica TCS SP8 microscope with an HC PL APO 93x/1.3 NA objective. Calcofluor was excited by UV at 405 nm, with emission detected between 420-470 nm.

CESA imaging was done on a Leica DMI6000 inverted microscope attached to a Yokogawa spinning disk confocal head with a 100x/1.4 NA objective. GFP was excited at 488 nm with a 525/50 nm emission filter. tdTomato was excited at 561 nm with a 593/40 nm emission filter. Excitation energy was adjusted to 4 mW power and exposure time was 200 ms. Z-stacks were obtained with a step size of 0.27 µm. Time-lapse images under the control or 1 M sorbitol condition (Figure S3B, Video S3) were recorded at a single focal plane at the plasma membrane with an interval of 10 s and a duration of 5 min. Time-lapse images of CSC relocalization before and after 1 M sorbitol treatment (Figure S3C, Video S5) were recorded at a single focal plane at the plasma membrane with an interval of 1 min and a duration of 15 min.

For Alexa568-tdCBM3a staining, five-day-old seedlings expressing GFP-CESA3 were stained with 200 nM Alexa568-tdCBM3a diluted in MS liquid in the dark for 30 min, washed with MS liquid, and imaged on the spinning disk confocal microscope described above with a 100x/1.4 NA objective. Alexa568 was excited at 561 nm with a 593/40 nm emission filter.

REM imaging was performed using a Leica TCS SP8 microscope with an HC PL APO 93x/1.3 NA objective. YFP excitation and emission parameters were consistent with those described above. For time-lapse imaging of REM nanodomain formation, a five-day-old YFP-REM1.2 seedling was placed under a 1X MS + 1% sucrose, 0.7% gelzan pad in a Nunc Lab-Tek chambered coverglass (Thermo Fisher, catalog #155360). A 0.5-cm diameter hole was created adjacent to the root tip. Time-lapse z-stack images were taken at a step-size of 0.3 µm, beginning at 0 min (pre-treatment) and then every minute for 10 min following the addition of 200 µL of 0.5 M sorbitol into the hole. To image REM nanodomain disappearance upon relief from hyperosmotic stress, a five-day-old YFP-REM1.2 seedling was pre-treated with 0.5 M sorbitol for 5 min and then placed under a 1X MS + 1% sucrose + 0.5 M sorbitol, 0.7% gelzan pad in a Nunc Lab-Tek chambered coverglass. Time-lapse z-stack imaging was performed under the same parameters as described above, except that 200 µL of 1X MS + 1% sucrose liquid solution was added to the hole instead of 0.5 M sorbitol.

### Image Analysis

3D renderings of confocal images were generated using Leica LAS X or AIVIA 15.0.0 software (Leica Microsystems). Hechtian structures were segmented in AIVIA. Quantitative image analyses were performed with ImageJ and Imaris 10.1.1 (Oxford Instruments).

CESA density was quantified in ImageJ. Z-stack images were first background-subtracted using the sliding paraboloid algorithm with a rolling ball radius of 30 pixels, and contrast was enhanced by setting saturated pixels to 0.35%. For each root cell, a substack containing CESA particles at the plasma membrane was generated and a maximum intensity projection was created. A region of interest (ROI) was defined using the polygon tool to exclude the Golgi, and its area was measured. Within this ROI, CESA particles were identified using the Find Maxima tool. The noise tolerance was set high enough to ensure that no maxima were detected in the background. The same noise tolerance threshold was consistently applied when comparing different conditions or genotypes in images acquired on the same day.

Time-lapse images of REM nanodomain formation and disappearance were processed in ImageJ by generating maximum projections of z-stacks at each time point using ImageJ. The resulting projections were concatenated and aligned using the StackReg plugin to correct for frame shifts.

To measure REM nanodomain density, z-stack images were similarly contrast-enhanced in ImageJ with saturated pixels set at 0.35%. A substack containing REM nanodomains at the plasma membrane was generated for each root cell, followed by maximum intensity projection and ROI selection using the polygon tool. The projection was then cropped and imported into Imaris, where REM nanodomains were detected using a 0.4-µm diameter threshold.

Colocalization analysis for dual marker lines was performed in ImageJ using the Coloc 2 plugin. For each root cell, an ROI was added to the ROI Manager and selected. Manders’ overlap coefficient was calculated using the Costes threshold regression method with 100 Costes randomizations.

### Biotin Treatment

One gram of five-day-old whole seedlings grown on MS plates was harvested into a nylon biopsy bag (Fisher Scientific, catalog #50-303-41). The bag was submerged in either 50 µM biotin in MS liquid medium (control) or 50 µM biotin in MS liquid supplemented with 0.5 M sorbitol for 15 minutes. After treatment, samples were washed with the corresponding solution (MS or MS + 0.5 M sorbitol) and gently blotted on a paper towel to remove excess liquid.

Samples were then flash-frozen in liquid nitrogen, ground into fine powder using a mortar and a pestle, and stored at −80°C for further use.

### Immunoblotting

To confirm biotinylation, 100 µl of finely ground frozen tissue powder from the biotin treatment was resuspended in 100 µl of 2× Laemmli buffer containing 120 mM Tris (pH 6.8), 4% SDS, 20% glycerol, 5% β-mercaptoethanol, and 0.05% bromophenol blue. Samples were boiled at 95°C for 5 minutes, and 10 µl of the supernatant was loaded onto a 10% SDS-PAGE gel.

Proteins were transferred to a 0.2 µm PVDF membrane (Bio-Rad, catalog #1704156) using the Trans-Blot Turbo Transfer System (Bio-Rad). Membranes were either stained with Coomassie Brilliant Blue R-250 (Bio-Rad, catalog #1610400) for 30 seconds and destained with a solution containing 40% methanol and 10% acetic acid, or blocked in 5% BSA in TBS and probed with HRP-conjugated streptavidin (Thermo Fisher, catalog #S911) diluted 1:5,000 in 5% BSA in TBST for 1 hour at room temperature. Chemiluminescence was detected using the Clarity ECL Western Substrate (Bio-Rad, catalog #1705061) and visualized with the ChemiDoc MP Imaging System (Bio-Rad).

### Protein Affinity Purification

One gram of the finely ground frozen tissue powder from the biotin treatment was resuspended in 2 mL of protein extraction buffer containing 50 mM Tris-HCl (pH 7.5), 150 mM NaCl, 0.1% SDS, 1% Triton X-100, 0.5% sodium deoxycholate, 1 mM EGTA, 1 mM DTT, 1× cOmplete protease inhibitor cocktail (Roche, catalog #COUEDTAF-RO), and 1 mM PMSF. The suspension was vortexed and incubated on a rotor wheel at 4°C for 10 min. Samples were then sonicated in an ice bath using a Branson 450 sonifier at 20% amplitude with 10 s on/10 s off pulses for a total of 2 min. Following sonication, samples were centrifuged at 12,000 g for 10 min at 4°C. A total of 2.5 mL of the resulting supernatant was loaded onto PD-10 desalting columns (Cytiva, catalog #17-0851-01) pre-equilibrated with equilibration buffer composed of 50 mM Tris-HCl (pH 7.5), 150 mM NaCl, 0.1% SDS, 1% Triton X-100, 0.5% sodium deoxycholate, 1 mM EGTA, and 1 mM DTT. Proteins were eluted with 3.5 mL of equilibration buffer into 5 mL LoBind tubes (Eppendorf, catalog #0030108302). For streptavidin pulldown, 200 µL of Dynabeads MyOne Streptavidin T1 beads (Thermo Fisher, catalog #65602) were washed with extraction buffer and then incubated with the desalted protein extract supplemented with 1× cOmplete protease inhibitor cocktail and 1 mM PMSF. The mixture was incubated overnight at 4°C on a rotor wheel. The following day, beads were separated using a magnetic rack, transferred to 1.5 mL LoBind tubes (Eppendorf, catalog #022431081), and subjected to a series of washes: twice with cold extraction buffer, twice with cold equilibration buffer, once with cold 1 M KCl, transferred to a fresh 1.5 mL LoBind tube, and washed twice more with cold extraction buffer. After the final wash, beads were separated on a magnetic rack to remove residual buffer and stored at −80°C for further use.

### Mass Spectrometry Sample Preparation

Frozen streptavidin beads were thawed and washed once with PBS prepared in HPLC-grade water (Thermo Fisher, catalog #022934-k7). The beads were transferred to a new 1.5 mL LoBind tube and washed five additional times with PBS in HPLC-grade water. After the final wash, beads were separated on a magnetic rack, and the buffer was carefully removed. Beads were then incubated with 80 µL trypsin digestion buffer containing 50 mM Tris (pH 7.5), 1 M Urea, 4 mM DTT, and 0.4 µg Trypsin at 25°C for 3 h with shaking at 1,200 rpm. Following incubation, beads were magnetically separated and washed twice with 60 µL of 1 M urea in 50 mM Tris (pH 7.5). The resulting supernatants were pooled and treated with 10 mM iodoacetamide at 25°C for 45 min in the dark with shaking at 1,200 rpm to alkylate cysteines. Digestion was continued by adding 0.5 µg of additional trypsin, followed by overnight incubation at 25°C with shaking at 1,200 rpm. The next day, a final 0.5 µg of trypsin was added, and samples were incubated for an additional 4 h at 25°C with shaking. Following digestion, 1% formic acid was added to acidify the samples, which were then vortexed and desalted using OMIX C18 pipette tips (10-100 µL, Agilent, catalog #A57003100) according to the manufacturer’s instructions. Peptides were dried using a speed vacuum centrifuge and resuspended in 0.1% formic acid for LC-MS/MS analysis. Samples were analyzed using a Q-Exactive HF hybrid quadrupole-Orbitrap mass spectrometer (Thermo Fisher), coupled to an Easy-nLC 1200 ultra-performance liquid chromatography system (Thermo Fisher).^30^

### Mass Spectrometry Data Analysis

Mass spectrometry data were analyzed using MaxQuant (version 2.4.2.0)^76^ and Perseus (version 2.0.10.0)^77^. For plasma membrane proteome comparisons between wild type and *cesa3^je^*^5^, raw data from TbID-YFP-LTI6b/Col-0, Col-0, TbID-YFP-LTI6b/*cesa3^je^*^5^, and *cesa3^je^*^5^ were searched together in MaxQuant using default parameters with the following modifications. The LFQ minimum ratio count was set to 2. Peptides were searched against the TAIR10 protein database (“TAIR10_pep_20101214”; https://www.arabidopsis.org/download/list?dir=Proteins%2FTAIR10_protein_lists), supplemented with a custom list of potential contaminants, such as trypsin, human keratin, streptavidin, BSA, YFP, TbID-YFP, along with the standard MaxQuant contaminant list. “Match between runs” was enabled with a match time window of 0.4 min and an alignment time window of 20 min. The resulting “proteinGroups.txt” file was imported into Perseus with LFQ intensities as the Main category. Proteins marked with reverse, contaminant, and only identified by site identifications were removed. Data were log2-transformed, grouped, and filtered to retain proteins with at least two valid values in at least one group. Data were imputed by replacing missing LFQ values from normal distribution using the total matrix mode with a width of 0.3 and down shift of 1.8. To remove all unspecific background labeling, each TbID-YFP-LTI6b-tagged group was compared to its respective genetic background by a one-sided Welch’s *t*-test with S0 of 0.5, FDR of 0.05, and 250 randomizations. The resulting filtered matrix was then used to compare TbID-YFP-LTI6b/*cesa3^je^*^5^ and TbID-YFP-LTI6b/Col-0 using a two-sided Student’s *t*-test with S0 of 0.5, FDR of 0.05, and 250 randomizations.

The same workflow was applied to the plasma membrane proteome comparison between the wild type and *rhm1-1*, using raw data of TbID-YFP-LTI6b/Col-0, Col-0, TbID-YFP-LTI6b/*rhm1-1*, and *rhm1-1* samples.

For REM1.2 proximity labeling experiments, raw data from TbID-YFP-REM1.2/Col-0 (control condition), TbID-YFP-LTI6b/Col-0 (control condition), Col-0 (control condition), TbID-YFP-REM1.2/Col-0 (0.5 M sorbitol), TbID-YFP-LTI6b/Col-0 (0.5 M sorbitol), and Col-0 (0.5 M sorbitol) were analyzed together using the same MaxQuant and Perseus workflow described above. To remove background labeling, TbID-YFP-REM1.2/Col-0 was compared to Col-0 under each condition by a one-sided Welch’s *t*-test with S0 of 0.5, FDR of 0.05, and 250 randomizations. The resulting filtered matrix was then used to compare the 0.5 M sorbitol treatment and the control condition in TbID-YFP-REM1.2/Col-0 using a two-sided Student’s *t*-test with S0 of 0.5, FDR of 0.05, and 250 randomizations.

The mass spectrometry proteomics data have been deposited to the ProteomeXchange Consortium (http://proteomecentral.proteomexchange.org) via the PRIDE^80^ partner repository with the dataset identifier PXD065856, PXD065860, and PXD065862.

### Transient Expression in Tobacco

*Nicotiana benthamiana* seeds were sown directly in PRO-MIX HP soil and grown in a growth chamber at 25°C under a 12-hour light/12-hour dark photoperiod. DNA constructs were transformed into *Agrobacterium tumefaciens* GV3101 (pMP90 and pSoup) strain using electroporation. Transformed Agrobacterium were cultured in 5 mL LB medium supplemented with antibiotics for two days, then pelleted by centrifugation at 3,000 g for 10 minutes. Pellets were resuspended and diluted in an infiltration medium containing 10 mM MgCl_2_, 10 mM MES to an OD600 of 0.5. Cultures were incubated at room temperature for 3 h with 150 µM acetosyringone. For infiltration, cultures were mixed with the Agrobacterium p19 strain at an OD600 of 0.2 and infiltrated to the abaxial side of leaves from 5-to 6-week-old Nicotiana plants. Infiltrated plants were kept in the growth chamber for an additional 3-5 days before imaging.

### FRET-FLIM

FRET-FLIM experiments in Nicotiana leaves were performed using a Leica TCS SP8 confocal microscope equipped with an HC PL APO 93x/1.3 NA objective. Leaves infiltrated with YFP-REM1.2 alone served as donor-only controls, while those co-infiltrated with YFP-REM1.2 and an mRFP-tagged construct served as donor + acceptor samples. Samples were incubated in either MS liquid medium or 1 M sorbitol to induce hyperosmotic stress. The donor (YFP) was excited at 514 nm using a pulsed white light laser operating at 80 MHz. Emission was collected between 525–570 nm using a HyD SMD hybrid detector. Laser power was adjusted to ensure a maximum of approximately one photon per laser pulse, and frame repetition was set to 20.

YFP fluorescence lifetimes were measured using the LAS X software. Under control conditions, ROIs were defined as narrow rectangular areas along the plasma membrane at cell borders. Under hyperosmotic stress, ROIs were selected based on individual YFP-REM1.2 nanodomains. Approximately 3-5 ROIs were analyzed per image. YFP lifetimes were calculated using an n-Exponential Reconvolution model with one component.

### Root vitrification for cryo-FIB milling and cryoET imaging

Five-day-old Arabidopsis seedlings expressing GFP-LTI6b grown on standard 1X MS media were treated with either fresh MS liquid or MS liquid supplemented with 1 M sorbitol for 5-10 min. Root tips were excised and transferred onto custom EM grids. Custom copper grids were fabricated from 100-µm-thick Cu foil using a Fablight FL4500 laser cutter. The grids were 3-mm-diameter discs containing a central slot (140 µm width, 1-2 mm long) designed to accommodate the thickness of Arabidopsis root tip. Grids were assembled into polished HPF planchettes (Leica) according to the waffle method^81^ and vitrified by high-pressure freezing using a Leica ICE high-pressure freezing platform. Following freezing, grids were clipped into Autogrid cartridges (Thermo Fisher Scientific) and stored in liquid nitrogen until subsequent cryo-focused ion beam (cryo-FIB) milling.

### FIB-milling

Following vitrification, the HPF frozen grids were kept in cryogenic temperature for all subsequent steps. First, they were loaded into a cryoFIB/SEM system Aquilos (Thermo Fisher Scientific) and regions of interest were localized using an integrated fluorescence microscope (iFLM Thermo Fisher Scientific). Lamellae were prepared using the serialized on-grid lift-in sectioning for tomography (SOLIST) method as described by Nguyen et al. (2024).^82^ Briefly, sample chunks were extracted using the EasyLift micromanipulator system (Thermo Fisher Scientific) with a Gallium ion source at 30 kV, divided into serial sections, and sequentially attached to all-gold support grids (UltrAuFoil, Quantifoil) using microsutures. Sections were secured to the gold foil by FIB-deposited microwelds and coated with organometallic platinum via GIS. Rough milling was performed at 1-0.5 nA to achieve a thickness of approximately 1-2 µm, followed by thinning and polishing at 100-30 pA to a final thickness of 170-220 nm. Lamella thickness and platinum layer integrity were monitored during final thinning using electron beam imaging at 2 kV and 13 pA.

### TEM data acquisition

Prepared lamellae were loaded under cryogenic temperature to a Krios G2 (Thermo Fisher Scientific) with a direct detector Falcon 4i (Thermo Fisher Scientific) and equipped with an energy filter Selectris X (Thermo Fisher Scientific) set to 10 eV slit for collection. Tomography 5 software (Thermo Fisher Scientific) was used to collect tomographic tilt-series in the .eer format in a dose-symmetric tilt scheme, ranging from +/-50 degrees in 2.5 degrees tilt step increments, accruing approximately 140 e^-^/Å^2^. The pixel size was 2.461Å and the defocus values ranged from 3.5 to 5.5 µm.

### Tomogram reconstruction and segmentation

The raw data in the .eer format was converted to .tiff file format using Relion3.^72^ Saved .tiff files were motion corrected with the MotionCor2,^73^ frames were re-aligned and 3D reconstructed with the AreTomo2 software,^74^ and the even/odd volumes were used for model training in CryoCare.^75^ The resulting trained model was used for denoising and merging the final 3D tomographic volume. The final volume was used for segmentation in ORS Dragonfly (version 2024.1). 4-7 representative slices from the tomogram were annotated for features of interest (ribosomes, plasma-membrane, cell wall) and used to train a U-Net neural network. U-net produced segmentations were then manually cleaned to remove artefacts (i.e. on the tomogram edges or due to ice).

### Quantification and Statistical Analysis

Grouped and scattered column graphs displaying mean ± SD were generated using Prism 10 (GraphPad). Histograms were created in Microsoft Excel, and volcano plots were generated using R. Statistical analyses were performed in Prism 10. Details of statistical tests, including sample sizes and significance levels, are provided in the figures and corresponding legends.

## Supplemental Video and able Titles and Legends

Video S1. Related to Figure 1. 3D rendering and segmentation of Hechtian structures and cell wall-plasma membrane attachment sites. The video was generated in AIVIA.

Video S2. Related to Figure 1. 3D segmentation of a cell wall-plasma membrane attachment site from cryogenic electron tomography. Cyan and yellow denote cellulose microfibrils and the membrane, respectively. See also Figure S1A.

Video S3. Related to Figure 4. GFP-CESA3 particle dynamics at the plasma membrane in root cells of 5-d-old seedlings under control conditions or after treatment with 1 M sorbitol. Scale bar: 5 µm. See also Figure S3B.

Video S4. Related to Figure 4. 3D rendering of a root cell of 5-d-old seedlings expressing GFP-LTI6b and tdTomato-CESA6 after treatment with 1 M sorbitol for 5 min. The video was generated in Imaris.

Video S5. Related to Figure 4. GFP-CESA3 particle dynamics at the plasma membrane in root cells of 5-d-old seedlings before and every 1 min after treatment with 1 M sorbitol. Scale bar: 5 µm. See also Figure S3C.

Video S6. Related to Figure 5. YFP-REM1.2 dynamics in root cells of 5-d-old seedlings before and every 1 min after treatment with 0.5 M sorbitol, or pre-stressed with 0.5 M sorbitol and every 1 min during recovery in MS medium. Scale bar: 10 µm.

Table S1. Tabs 1 and 2, proximity labeling results comparing plasma membrane proteome between *cesa3^je^*^5^ and wild type Col-0, related to Figure 3D and S2H. Tabs 3 and 4, proximity labeling results comparing plasma membrane proteome between *rhm1-1* and wild type Col-0, related to Figure 3E and S2I. Tabs 5 and 6, proximity labeling results of TbID-YFP-REM1.2 in the presence or absence of 0.5 M sorbitol, related to Figure 7A and 7B. Tab 7, list of primers used in this study, related to STAR Methods.

