## Supplemental figures for "Cellulose Synthase Complexes and Remorins Mediate Stress Resilience Through Cell Wall-Plasma Membrane Attachments"

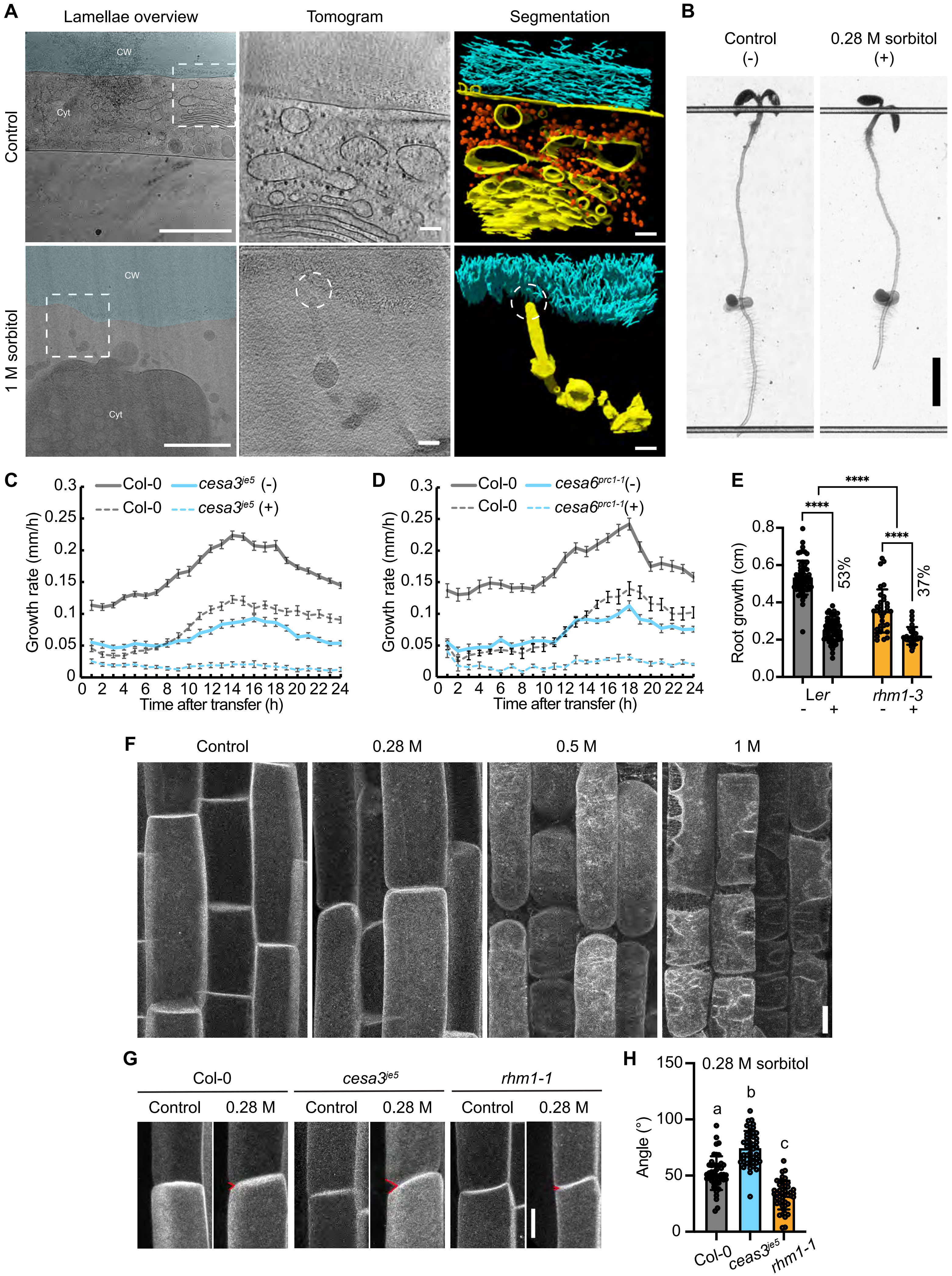

A
Lamellae overview
Tomogram
Segmentation
Control
1 M sorbitol
CW
Cyt
B
Control (-)
0.28 M sorbitol (+)
C
Col-0
cesa3je5 (-)
Col-0
cesa3je5 (+)
Growth rate (mm/h)
Time after transfer (h)
D
Col-0
cesa6prc1-1(-)
Col-0
cesa6prc1-1(+)
Growth rate (mm/h)
Time after transfer (h)
E
Root growth (cm)
53%
37%
Ler
rhm1-3
F
Control
0.28 M
0.5 M
1 M
G
Col-0
cesa3je5
rhm1-1
Control
0.28 M
H
0.28 M sorbitol
Angle (°)
a
b
c
Col-0
ceas3je5
rhm1-1

**Figure S1. Related to Figures 1, 2, and 3.**
(A) Lamellae overviews, tomograms, and corresponding 3D segmentations of root cells under control conditions or following treatment with 1 M sorbitol. In the lamellae overviews, white dashed boxes mark regions of interest shown at higher magnification in the tomograms and 3D segmentations. CW, cell wall; Cyt, cytoplasm. Cell walls are shaded in light cyan. Tomograms show projections from five consecutive slices. In the segmentations, cyan, yellow, and orange indicate cellulose microfibrils in the cell wall, membrane, and ribosomes, respectively. White dashed circles highlight wall-membrane attachment sites. Scale bars: 1 μm (lamellae overviews) and 100 nm (tomograms and segmentations).
(B) 5-d-old Col-0 seedlings imaged 1 day after transfer to either fresh MS media (-) or MS media supplemented with 0.28 M sorbitol (+). Black dots indicate the position of the root tip at the time of transfer. Scale bar: 0.2 cm.
(C) Measurements of root growth rate in Col-0 and *cesa3^je5^* seedlings immediately after transfer to either fresh MS media (-) or MS media supplemented with 0.28 M sorbitol (+) and monitored for 24 h. Error bars indicate SE. $n \geq 11$ seedlings per genotype per treatment.
(D) Measurements of root growth rate in Col-0 and *cesa6^prc1-1^* seedlings immediately after transfer to either fresh MS media (-) or MS media supplemented with 0.28 M sorbitol (+) and monitored for 24 h. Error bars indicate SE. $n \geq 8$ seedlings per genotype per treatment.
(E) Root growth of wild type and *rhm1-3* mutants 1 day after transfer to either fresh MS media (-) or MS media supplemented with 0.28 M sorbitol (+). $n \geq 30$ seedlings per genotype per treatment. Error bars indicate SD. ****$p < 0.0001$ by two-way ANOVA.
(F) Laser scanning confocal images of root epidermal cells in 5-d-old YFP-LTI6b seedlings under the control condition or treated with 0.28 M, 0.5 M, or 1 M sorbitol, respectively, for 5 min. Scale bar: 10 μm.
(G) Laser scanning confocal images of root epidermal cells treated with control or 0.28 M sorbitol for 5 min in 5-d-old seedlings expressing YFP-LTI6b in Col-0, *cesa3^je5^*, or *rhm1-1* background, respectively. Red carets indicate angles measured in (H). Scale bar: 10 μm.
(H) Measurements of the angle between two cell corners in Col-0, *cesa3^je5^*, and *rhm1-1* following treatment with 0.28 M sorbitol for 5 min. $n \geq 45$ cells from at least 9 seedlings per genotype. Error bars indicate SD. Different letters indicate significant differences by one-way ANOVA and Tukey's test.

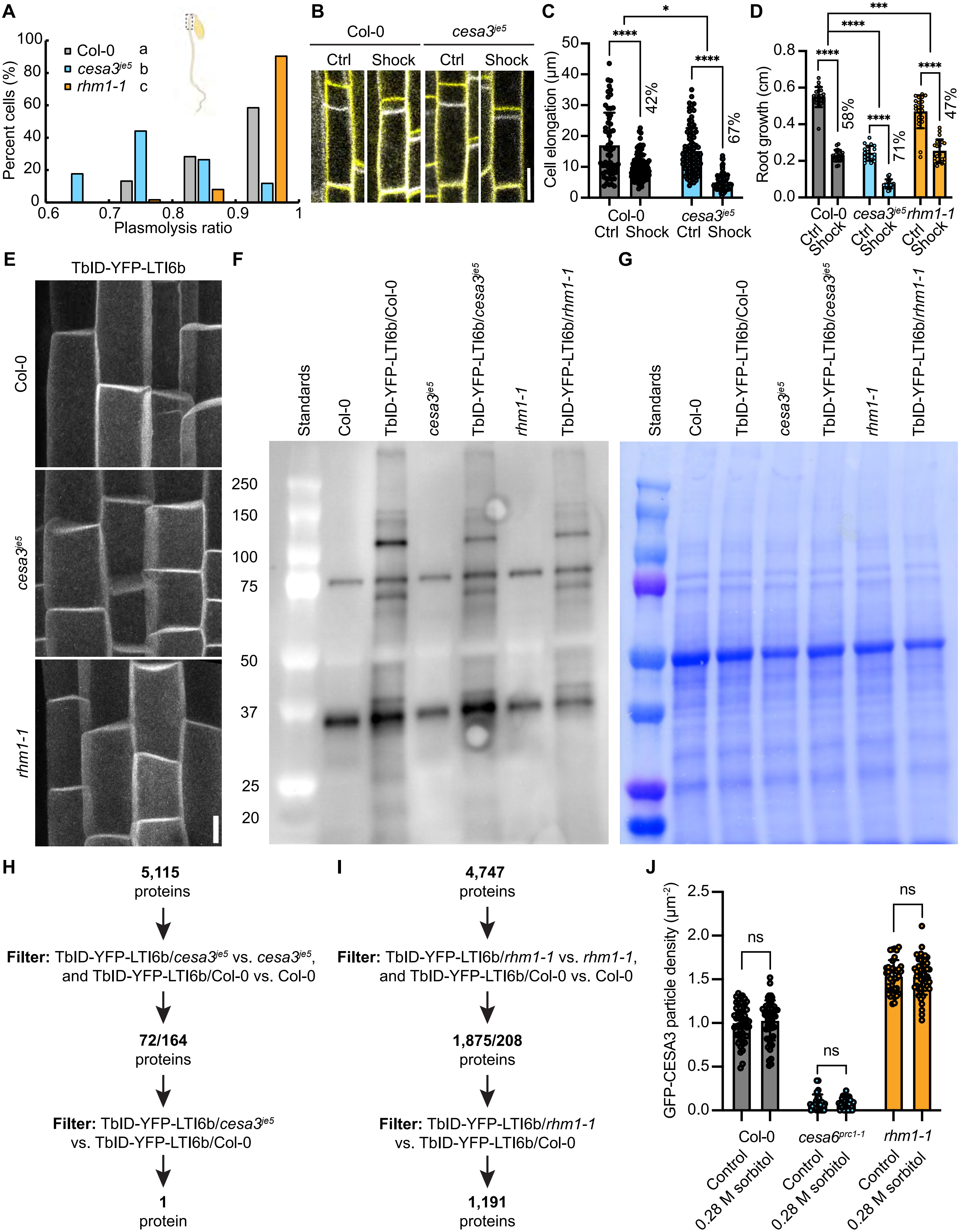

A
Col-0 a
cesa3je5 b
rhm1-1 c
Percent cells (%)
Plasmolysis ratio
B
Col-0
cesa3je5
Ctrl
Shock
C
Cell elongation (µm)
42%
67%
Col-0
cesa3je5
Ctrl Shock Ctrl Shock
D
Root growth (cm)
58%
71%
47%
Col-0
cesa3je5
rhm1-1
E
TbID-YFP-LTI6b
Col-0
cesa3je5
rhm1-1
F
Standards
Col-0
TbID-YFP-LTI6b/Col-0
cesa3je5
TbID-YFP-LTI6b/cesa3je5
rhm1-1
TbID-YFP-LTI6b/rhm1-1
250
150
100
75
50
37
25
20
G
Standards
Col-0
TbID-YFP-LTI6b/Col-0
cesa3je5
TbID-YFP-LTI6b/cesa3je5
rhm1-1
TbID-YFP-LTI6b/rhm1-1
H
5,115 proteins
Filter: TbID-YFP-LTI6b/cesa3je5 vs. cesa3je5, and TbID-YFP-LTI6b/Col-0 vs. Col-0
72/164 proteins
Filter: TbID-YFP-LTI6b/cesa3je5 vs. TbID-YFP-LTI6b/Col-0
1 protein
I
4,747 proteins
Filter: TbID-YFP-LTI6b/rhm1-1 vs. rhm1-1, and TbID-YFP-LTI6b/Col-0 vs. Col-0
1,875/208 proteins
Filter: TbID-YFP-LTI6b/rhm1-1 vs. TbID-YFP-LTI6b/Col-0
1,191 proteins
J
GFP-CESA3 particle density (µm-2)
ns
Col-0
cesa6prc1-1
rhm1-1
Control
0.28 M sorbitol

**Figure S2. Related to Figure 3.**
(A) Histogram showing plasmolysis ratio (protoplast area to cell area) in hypocotyl cells of 3-d-old etiolated seedlings expressing YFP-LTI6b in Col-0, *cesa3$^{je5}$*, or *rhm1-1* background, respectively. Different letters indicate significant differences by one-way ANOVA and Tukey's test. $n \geq 34$ cells from at least 5 seedlings per genotype.
(B) Laser scanning confocal images of root epidermal cells of 5-d-old seedlings expressing YFP-LTI6b in Col-0 and *cesa3$^{je5}$*. Seedlings were treated under the control condition or with 0.5 M sorbitol for 5 min, then transferred back to MS + 1% sucrose medium. Cells within the same region of interest were imaged immediately after transfer (white) and again 1 hour later (yellow). Scale bar: 20 μm.
(C) Quantification of cell elongation in root epidermal cells of 5-d-old seedlings expressing YFP-LTI6b in Col-0 and *cesa3$^{je5}$* from (B). Cell elongation was calculated as the change in cell length between the two time points. Error bars indicate SD. $n \geq 61$ cells from at least 7 seedlings per genotype per treatment. *$p < 0.05$ and ****$p < 0.0001$ by two-way ANOVA.
(D) Root growth of wild type and *cesa3$^{je5}$* 1 day after treatment under the control condition or with 0.5 M sorbitol for 30 min, then transferred back to MS + 1% sucrose medium. $n \geq 19$ seedlings per genotype per treatment. Error bars indicate SD. ****$p < 0.0001$ by two-way ANOVA.
(E) Laser scanning confocal images of root epidermal cells from 5-d-old seedlings of TbID-YFP-LTI6b/Col-0, TbID-YFP-LTI6b/*cesa3$^{je5}$*, and TbID-YFP-LTI6b/*rhm1-1*, respectively. Scale bar: 10 μm.
(F) Validation of TbID activity in 5-d-old seedlings treated with 50 μM biotin for 15 min assessed by immunoblotting using HRP-conjugated streptavidin.
(G) Coomassie Brilliant Blue-stained membrane showing the loading controls in (F).
(H) Workflow of filtering steps to identify differentially enriched proteins in TbID-YFP-LTI6b/*cesa3$^{je5}$* compared to TbID-YFP-LTI6b/Col-0.
(I) Workflow of filtering steps to identify differentially enriched proteins in TbID-YFP-LTI6b/*rhm1-1* compared to TbID-YFP-LTI6b/Col-0.
(J) Quantification of GFP-CESA3 particle density at the plasma membrane in root epidermal cells of 5-d-old seedlings of Col-0, *cesa6$^{prc1-1}$*, and *rhm1-1* under control conditions or after treatment with 0.28 M sorbitol for 5 minutes. Error bars indicate SD. $n \geq 25$ cells from at least 6 seedlings per genotype per treatment. ns, no significance by Student's *t*-test.

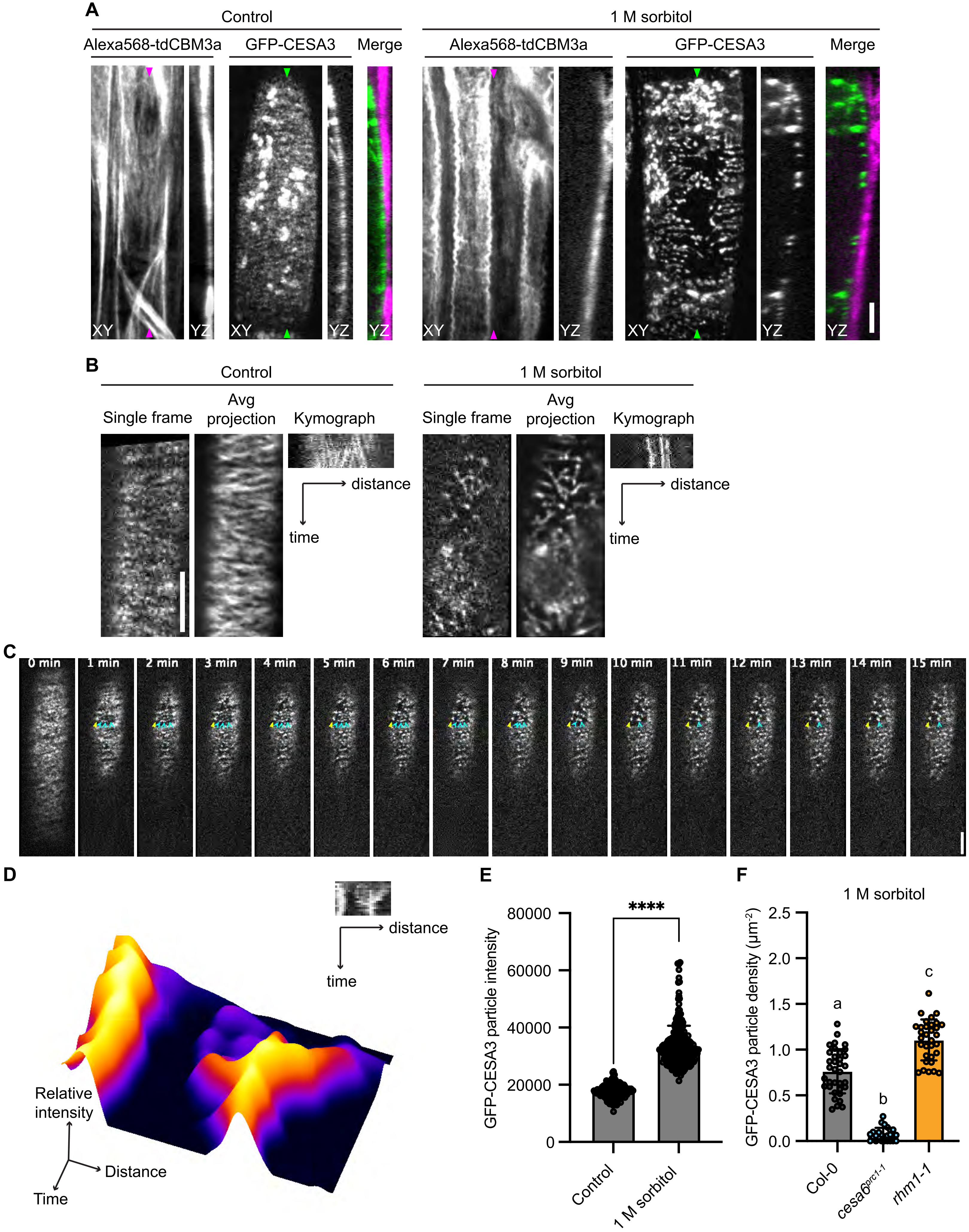
A
Control
Alexa568-tdCBM3a
GFP-CESA3
Merge
1 M sorbitol
XY
YZ
B
Single frame
Avg projection
Kymograph
distance
time
C
0 min
1 min
2 min
3 min
4 min
5 min
6 min
7 min
8 min
9 min
10 min
11 min
12 min
13 min
14 min
15 min
D
Relative intensity
Distance
Time
E
GFP-CESA3 particle intensity
****
80000
60000
40000
20000
0
Control
1 M sorbitol
F
GFP-CESA3 particle density (μm⁻²)
2.5
2.0
1.5
1.0
0.5
0.0
a
b
c
Col-0
cesa6prc1-1
rhm1-1

**Figure S3. Related to Figure 4.**
(A) Spinning disk confocal images of root epidermal cells in 5-d-old seedlings expressing GFP-CESA3 co-stained with 200 nM Alexa568-tdCBM3a under the control condition or treated with 1 M sorbitol for 5 min. XY and YZ projections are shown. The positions where YZ projections were made are indicated by arrowheads. Scale bar: 5 μm.
(B) Spinning disk confocal images showing GFP-CESA3 particle motility in root epidermal cells of 5-d-old seedlings under control conditions or after treatment with 1 M sorbitol for 5 minutes. Scale bar: 5 μm.
(C) Time-series spinning disk confocal images showing GFP-CESA3 particle dynamics at the same single plane before (0 min) and every 1 min after treatment with 1 M sorbitol for a total of 15 minutes. Scale bar: 5 μm. Yellow arrowhead points to a CSC cluster present immediately after the treatment. Cyan arrowheads point to multiple CSCs aggregating together.
(D) Surface plot and kymograph of the particles highlighted in yellow and cyan arrowheads in (C).
(E) Quantification of GFP-CESA3 particle intensity in root epidermal cells of 5-d-old seedlings under control conditions or after treatment with 1 M sorbitol for 5 minutes. Error bars indicate SD. $n \geq 156$ particles from at least 7 seedlings per treatment. ****$p < 0.0001$ by Student's *t*-test.
(F) Quantification of GFP-CESA3 particle density at the plasma membrane in root epidermal cells of 5-d-old seedlings of Col-0, *cesa6*$^{prc1\text{-}1}$, and *rhm1-1* after treatment with 1 M sorbitol for 5 minutes. Error bars indicate SD. Different letters indicate significant differences by one-way ANOVA and Tukey's test. $n \geq 24$ cells from 9 seedlings per genotype.

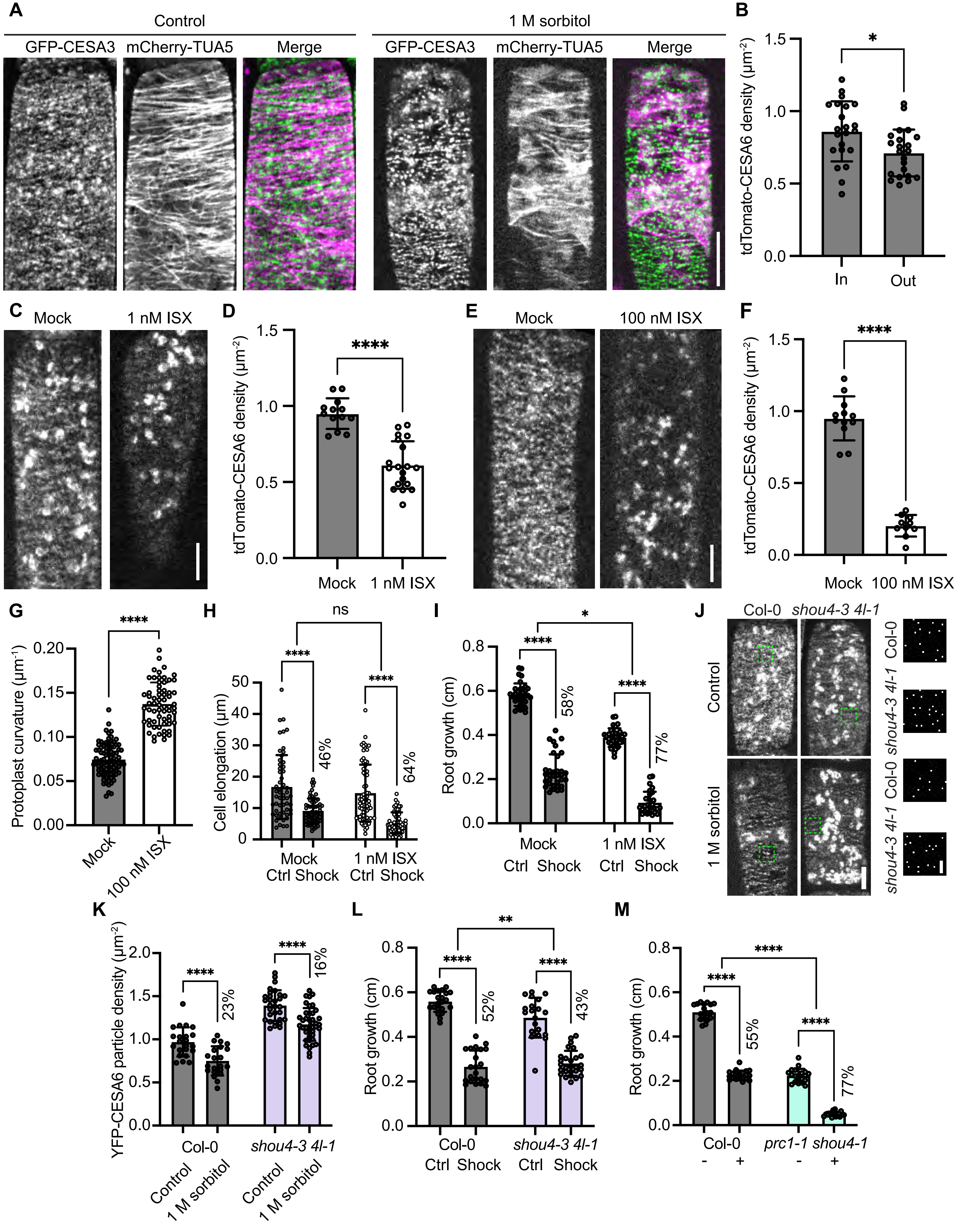

A
Control
1 M sorbitol
GFP-CESA3
mCherry-TUA5
Merge
B
tdTomato-CESA6 density (µm⁻²)
In
Out
C
Mock
1 nM ISX
D
1 nM ISX
E
100 nM ISX
F
G
Protoplast curvature (µm⁻¹)
H
Cell elongation (µm)
ns
46%
64%
Mock
Ctrl Shock
I
Root growth (cm)
58%
77%
J
Col-0
shou4-3 4l-1
K
YFP-CESA6 particle density (µm⁻²)
23%
16%
L
52%
43%
M
55%
prc1-1 shou4-1

**Figure S4. Related to Figure 4.**
(A) Spinning disk confocal images of root epidermal cells of 5-d-old seedlings expressing GFP-CESA3 and mCherry-TUA5 under control conditions or after treatment with 1 M sorbitol for 5 minutes. Scale bar: 10 μm.
(B) Quantification of tdTomato-CESA6 particle density at the plasma membrane in and out of regions associated with microtubules in 5-d-old seedlings treated with 1 M sorbitol for 5 minutes. $n = 22$ cells from 6 seedlings. Error bars indicate SD. *$p < 0.05$ by Student's *t*-test.
(C) Spinning disk confocal images of root epidermal cells in 5-d-old tdTomato-CESA6 seedlings treated with or without 1 nM ISX for 20 min. Scale bar: 5 μm.
(D) Quantification of tdTomato-CESA6 particle density at the plasma membrane from (C). $n \geq 13$ cells from at least 3 seedlings per treatment. Error bars indicate SD. ****$p < 0.0001$ by Student's *t*-test.
(E) Spinning disk confocal images of root epidermal cells in 5-d-old tdTomato-CESA6 seedlings treated with or without 100 nM ISX for 20 min. Scale bar: 5 μm.
(F) Quantification of tdTomato-CESA6 particle density at the plasma membrane from (E). $n \geq 19$ cells from at least 5 seedlings per treatment. Error bars indicate SD. ****$p < 0.0001$ by Student's *t*-test.
(G) Measurements of protoplast curvature in root epidermal cells of 5-d-old YFP-LTI6b seedlings treated with or without 100 nM ISX for 20 minutes. Plasmolysis was subsequently induced by treatment with 0.5 M sorbitol for 5 minutes. $n \geq 64$ cells from at least 5 seedlings per treatment. Error bars indicate SD. ****$p < 0.0001$ by Student's *t*-test.
(H) Quantification of cell elongation in root epidermal cells of 5-d-old Col-0 seedlings expressing YFP-LTI6b. 4-d-old seedlings were first transferred to MS medium in the presence or absence of 1 nM ISX and let growt for 1 day. Seedlings were then treated under the control condition or with 0.5 M sorbitol for 5 min, and transferred back to MS medium in the presence or absence of 1 nM ISX. Cells within the same region of interest were imaged immediately after transfer and again 1 hour later. Cell elongation was calculated as the change in cell length between the two time points. Error bars indicate SD. $n \geq 55$ cells from at least 5 seedlings per genotype per treatment. ns, no significance; ****$p < 0.0001$ by two-way ANOVA.
(I) Root growth of wild type 1 day after treatment under the control condition or with 0.5 M sorbitol for 30 min, then transferred back to MS medium in the presence or absence of 1 nM ISX. $n \geq 29$ seedlings per treatment. Error bars indicate SD. *$p < 0.05$ and ****$p < 0.0001$ by two-way ANOVA.
(J) Spinning disk confocal images of YFP-CESA6 in root epidermal cells of 5-d-old seedlings of Col-0 and *shou4-3 4l-1* under the control condition or induced by 1 M sorbitol for 5 min. Scale bar: 5 μm. Magnified views of the regions outlined by green dashed boxes, highlighting detected YFP-CESA6 particles, are shown on the right. Scale bar: 1 μm.
(K) Quantification of YFP-CESA6 particle density in root epidermal cells of 5-d-old seedlings of Col-0 and *shou4-3 4l-1* under control conditions or after treatment with 1 M sorbitol for 5 minutes. Error bars indicate SD. $n \geq 29$ cells from at least 6 seedlings per treatment. ****$p < 0.0001$ by Student's *t*-test.
(L) Root growth of wild type and *shou4-3 4l-1* 1 day after treatment under the control condition or with 0.5 M sorbitol for 30 min, then transferred back to MS + 1% sucrose medium. $n \geq 20$ seedlings per genotype per treatment. Error bars indicate SD. **$p < 0.01$ and ****$p < 0.0001$ by two-way ANOVA.
(M) Root growth of wild type and *prc1-1 shou4-1* mutants 1 day after transfer to either fresh MS media (-) or MS media supplemented with 0.28 M sorbitol (+). $n \geq 19$ seedlings per genotype per treatment. Error bars indicate SD. ****$p < 0.0001$ by two-way ANOVA.

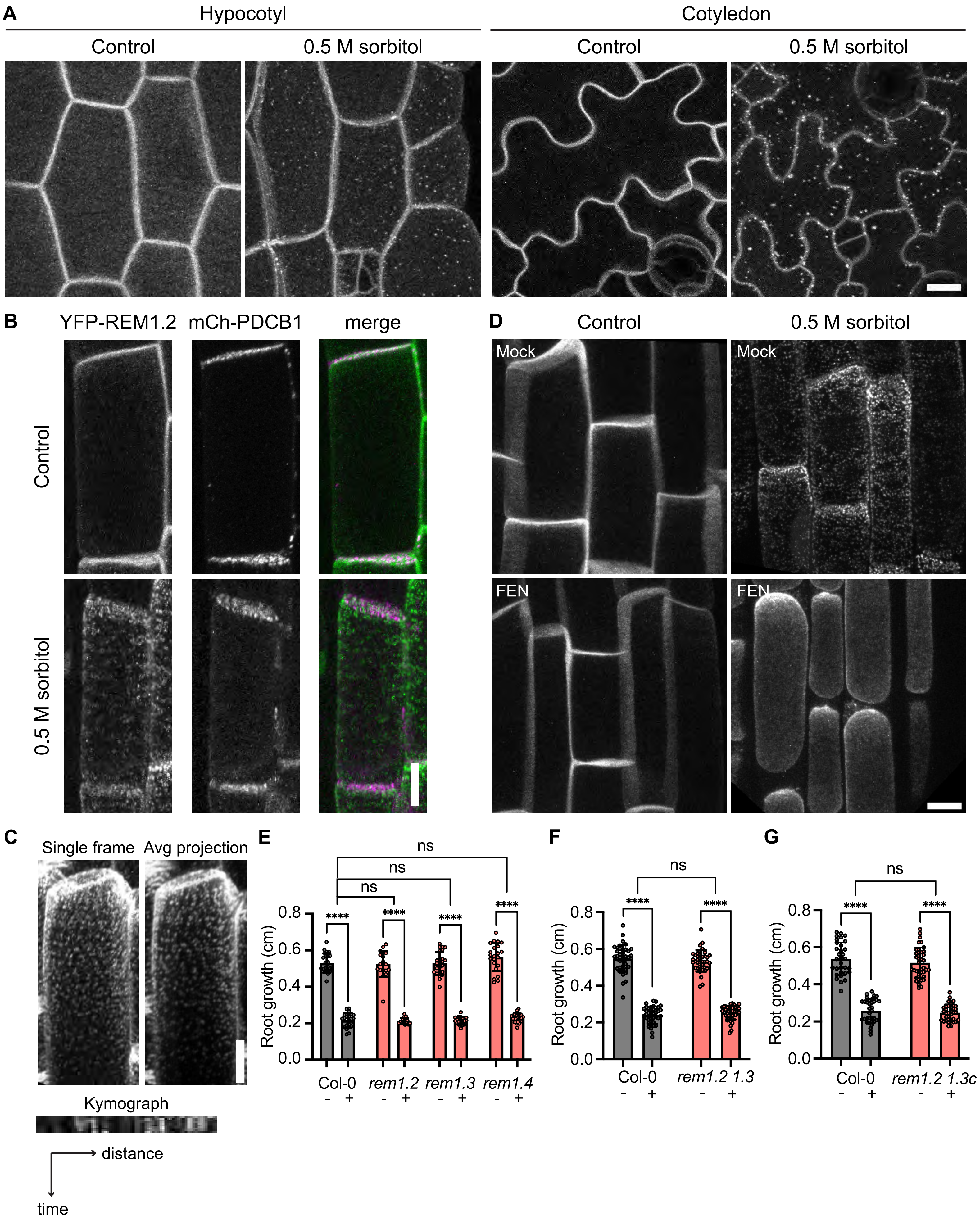

A
Hypocotyl
Cotyledon
Control
0.5 M sorbitol
Control
0.5 M sorbitol
B
YFP-REM1.2
mCh-PDCB1
merge
Control
0.5 M sorbitol
D
Control
0.5 M sorbitol
Mock
Mock
FEN
FEN
C
Single frame
Avg projection
Kymograph
distance
time
E
ns
ns
ns
****
****
****
****
Root growth (cm)
0.0
0.2
0.4
0.6
0.8
Col-0
rem1.2
rem1.3
rem1.4
- +
F
ns
****
****
Root growth (cm)
Col-0
rem1.2 1.3
- +
G
ns
****
****
Root growth (cm)
Col-0
rem1.2 1.3c
- +

**Figure S5. Related to Figure 5.**
(A) Laser scanning confocal images of hypocotyl and cotyledons cells of 5-d-old YFP-REM1.2 seedlings under the control condition or treated with 0.5 M sorbitol for 5 min. Scale bar: 10 μm.
(B) Laser scanning confocal images of root epidermal cells in 5-d-old seedlings expressing YFP-REM1.2 and mCherry-PDCB1 under the control condition or treated with 0.5 M sorbitol for 5 min. Scale bar: 10 μm.
(C) Laser scanning confocal images showing immobile YFP-REM1.2 nanodomains in root epidermal cells of 5-d-old seedlings upon treatment with 0.5 M sorbitol. Scale bar: 10 μm.
(D) Laser scanning confocal images of root epidermal cells in 5-d-old YFP-REM1.2 seedlings, taken 1 day after treatment with or without 100 μg/mL fenpropimorph (FEN), followed by a 5-min treatment with 0.5 M sorbitol. Scale bar: 10 μm.
(E) Root growth of wild type, *rem1.2*, *rem1.3*, and *rem1.4* single mutants 1 day after transfer to either fresh MS media (-) or MS media supplemented with 0.28 M sorbitol (+). $n \geq 17$ seedlings per genotype per treatment. Error bars indicate SD. ns, no significance; ****$p < 0.0001$ by two-way ANOVA.
(F) and (G) Root growth of wild type, *rem1.2 1.3*, and *rem1.2 1.3c* mutants 1 day after transfer to either fresh MS media (-) or MS media supplemented with 0.28 M sorbitol (+). $n \geq 30$ seedlings per genotype per treatment. Error bars indicate SD. ns, no significance; ****$p < 0.0001$ by two-way ANOVA.

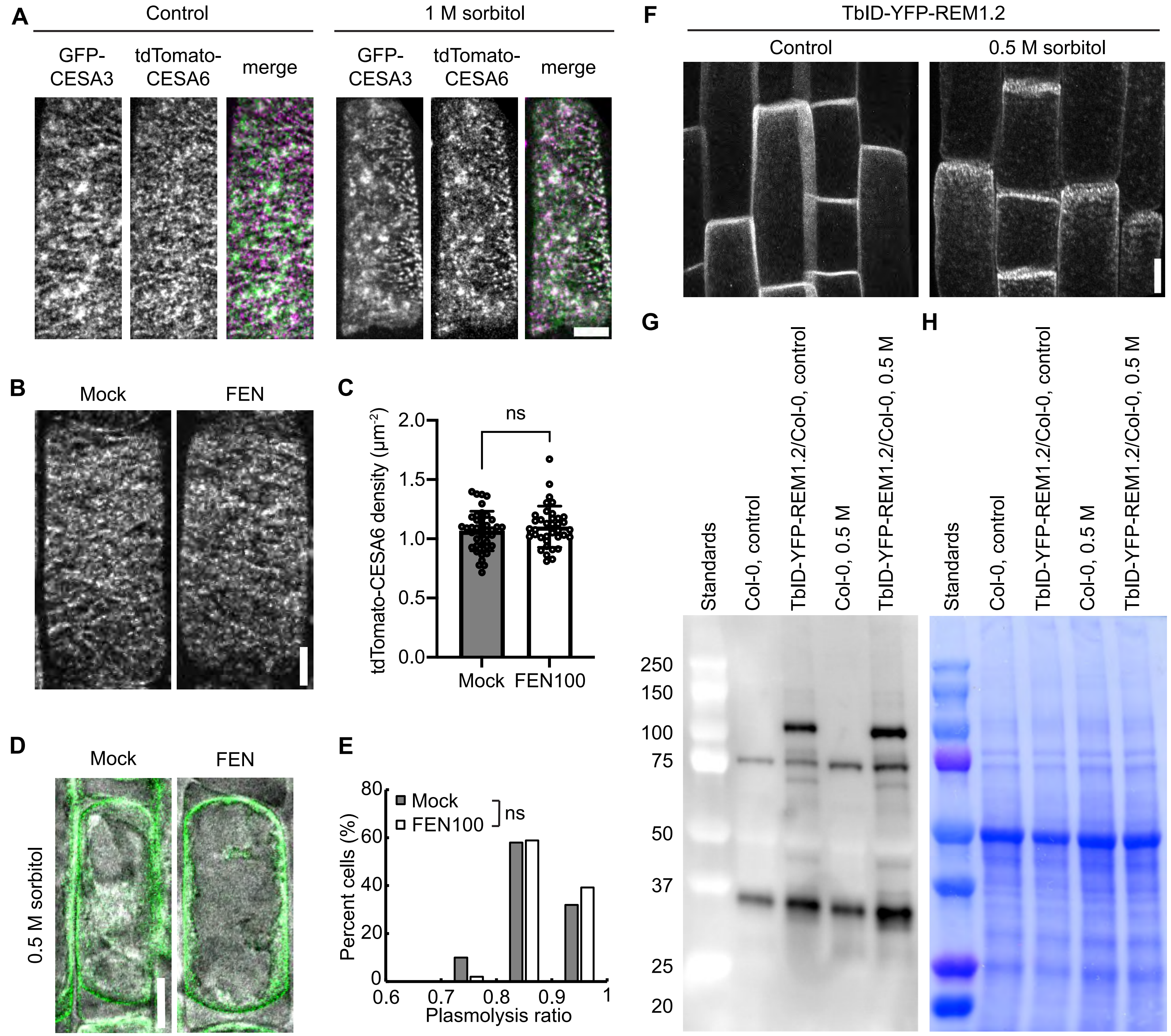


**Figure S6. Related to Figures 6 and 7.**

(A) Spinning disk confocal images of root epidermal cells in 5-d-old seedlings expressing GFP-CESA3 and tdTomato-CESA6 under the control condition or treated with 1 M sorbitol for 5 min. Scale bar: 5 μm.

(B) Spinning disk confocal images of root epidermal cells in 5-d-old GFP-CESA3 seedlings, taken 1 day after treatment with or without 100 μg/mL fenpropimorph (FEN). Scale bar: 5 μm.

(C) Quantification of GFP-CESA3 particle density at the plasma membrane from (B). Error bars indicate SD. ns, no significance by Student's *t*-test. $n \geq 37$ cells from at least 10 seedlings per treatment.

(D) Laser scanning confocal images of root epidermal cells in 5-d-old GFP-LTI6b seedlings, taken 1 day after treatment with or without 100 μg/mL FEN, followed by a 5-min treatment with 0.5 M sorbitol. Scale bar: 10 μm.

(E) Histogram showing plasmolysis ratio in root epidermal cells of 5-d-old GFP-LTI6b seedlings treated with or without 100 μg/mL FEN for 1 day. ns, no significance by Student's *t*-test. $n \geq 50$ cells from at least 5 seedlings per treatment.

(F) Laser scanning confocal images of root epidermal cells from 5-d-old seedlings of TbID-YFP-REM1.2 under the control condition or treated with 0.5 M sorbitol for 5 min. Scale bar: 10 μm.

(G) Validation of TbID activity in 5-d-old seedlings treated with 50 μM biotin for 15 min in the presence or absence of 0.5 M sorbitol assessed by immunoblotting using HRP-conjugated streptavidin.

(H) Coomassie Brilliant Blue-stained membrane showing the loading controls in (G).

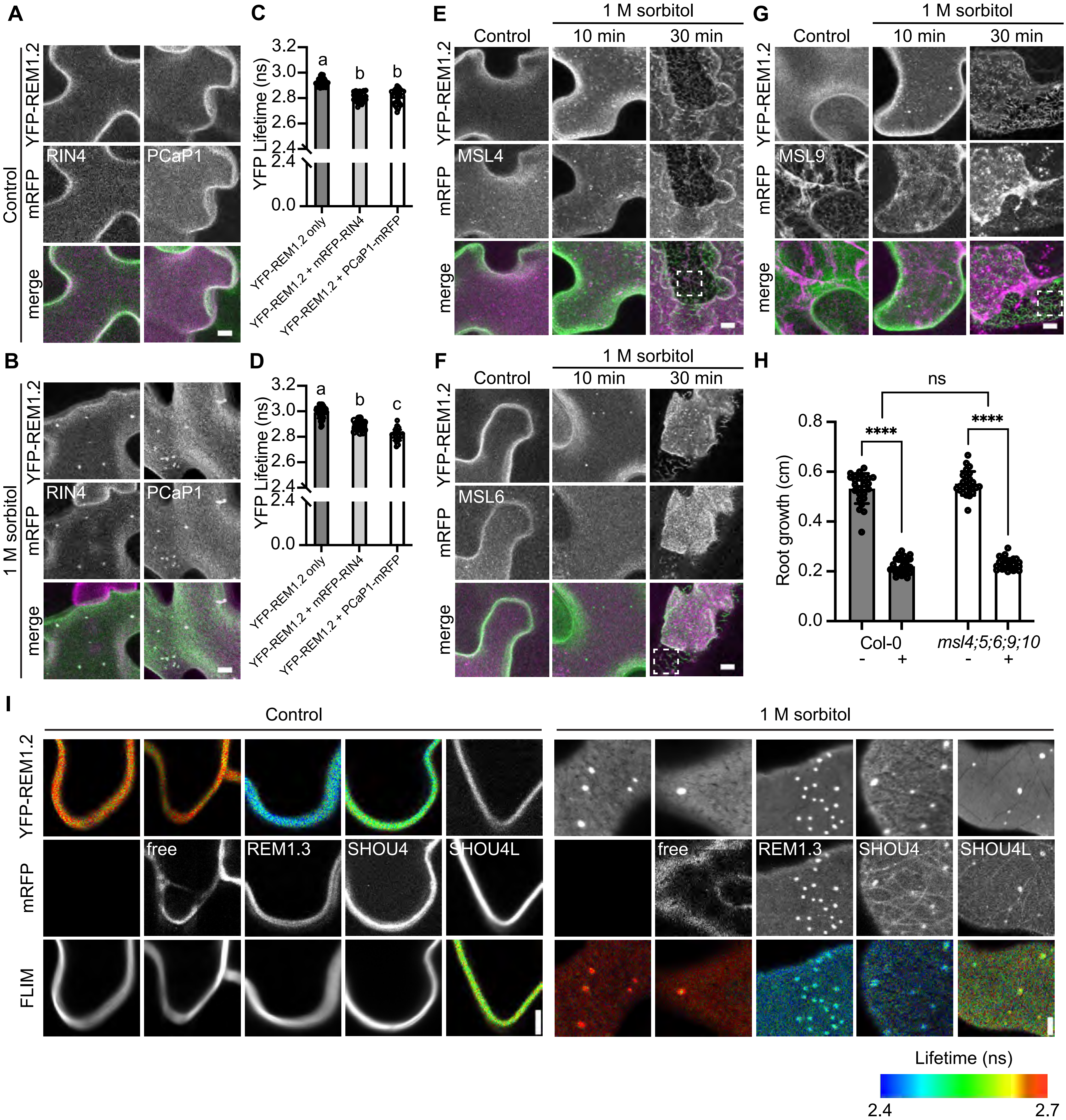

A
B
C
D
E
F
G
H
I
Control
1 M sorbitol
YFP-REM1.2
mRFP
merge
RIN4
PCaP1
YFP Lifetime (ns)
YFP-REM1.2 only
YFP-REM1.2 + mRFP-RIN4
YFP-REM1.2 + PCaP1-mRFP
10 min
30 min
MSL4
MSL6
MSL9
Root growth (cm)
Col-0
msl4;5;6;9;10
ns
****
free
REM1.3
SHOU4
SHOU4L
FLIM
Lifetime (ns)
2.4
2.7

**Figure S7. Related to Figure 7.**
(A) and (B) Laser scanning confocal images of tobacco leaf epidermal cells co-expressing YFP-REM1.2 with mRFP-RIN4 or PCaP1-mRFP under control conditions (A) or upon 1 M sorbitol treatment for 10 min (B). Scale bars: 5 μm.
(C) and (D) Measurements of YFP lifetime under control conditions (C) or upon 1 M sorbitol treatment for 10 min (D). Error bars indicate SD. Different letters indicate significant differences by one-way ANOVA and Tukey's test. $n \geq 25$ regions of interest from at least 3 infiltrated plants per construct combination.
(E-G) Laser scanning confocal images of tobacco leaf epidermal cells co-expressing YFP-REM1.2 with MSL4-mRFP (E), MSL6-mRFP (F), or MSL9-mRFP (G) under control conditions or treated with 1 M sorbitol for 10 min or 30 min. Examples of Hechtian reticulum are indicated by dashed boxes in the merged images. Scale bars: 5 μm.
(H) Root growth of wild type and *msl4;5;6;9;10* mutants 1 day after transfer to either fresh MS media (-) or MS media supplemented with 0.28 M sorbitol (+). $n \geq 24$ seedlings per genotype per treatment. Error bars indicate SD. ns, no significance; ****$p < 0.0001$ by two-way ANOVA.
(I) Fluorescence lifetime imaging microscopy (FLIM) in tobacco leaf epidermal cells expressing YFP-REM1.2 alone, or co-expressing YFP-REM1.2 with free mRFP, mRFP-REM1.3, SHOU4-mRFP, or SHOU4L-mRFP, respectively, under control conditions or treated with 1 M sorbitol for 10 min. Scale bars: 5 μm.
